# Multi-color fluorescence fluctuation spectroscopy in living cells via spectral detection

**DOI:** 10.1101/2020.12.18.423407

**Authors:** Valentin Dunsing, Annett Petrich, Salvatore Chiantia

## Abstract

Signaling pathways in biological systems rely on specific interactions between multiple biomolecules. Fluorescence fluctuation spectroscopy provides a powerful toolbox to quantify such interactions directly in living cells. Cross-correlation analysis of spectrally separated fluctuations provides information about inter-molecular interactions but is usually limited to two fluorophore species. Here, we present scanning fluorescence spectral correlation spectroscopy (SFSCS), a versatile approach that can be implemented on commercial confocal microscopes, allowing the investigation of interactions between multiple protein species at the plasma membrane. We demonstrate that SFSCS enables cross-talk-free cross-correlation, diffusion and oligomerization analysis of up to four protein species labeled with strongly overlapping fluorophores. As an example, we investigate the interactions of influenza A virus (IAV) matrix protein 2 with two cellular host factors simultaneously. We furthermore apply raster spectral image correlation spectroscopy for the simultaneous analysis of up to four species and determine the stoichiometry of ternary IAV polymerase complexes in the cell nucleus.

## INTRODUCTION

Living cells rely on transport and interaction of biomolecules to perform their diverse functions. To investigate the underlying molecular processes in the native cellular environment, minimally invasive techniques are needed. Fluorescence fluctuation spectroscopy (FFS) approaches provide a powerful toolbox that fulfills this aim (1–3). FFS takes advantage of inherent molecular dynamics present in biological systems, for example diffusion, to obtain molecular parameters from fluctuations of the signal emitted by an ensemble of fluorescent molecules. More in detail, the temporal evolution of such fluctuations allows the quantification of intracellular dynamics. In addition, concentration and oligomerization state of molecular complexes can be determined by analyzing the magnitude of fluctuations. Finally, hetero-interactions of different molecular species can be detected by cross-correlation analysis of fluctuations emitted by spectrally separated fluorophores (4). Over the last two decades, several experimental FFS schemes such as raster image (cross-) correlation spectroscopy (RI(C)CS) (5, 6), (cross-correlation) Number&Brightness analysis (7, 8), and imaging FCS (9) have been developed, extending the concept of traditional single-point fluorescence (cross-) correlation spectroscopy (F(C)CS) (10). A further interesting example of FFS analysis relevant in the field of cell biology is represented by scanning F(C)CS (SF(C)CS). Using a scanning path perpendicular to the plasma membrane (PM), this technique provides enhanced stability and the ability to probe slow membrane dynamics (11), protein interactions (12, 13) and oligomerization (14) at the PM of cells.

FFS studies are conventionally limited to the analysis of two spectrally distinguished species, due to i) broad emission spectra of fluorophores with consequent cross-talk artefacts, and ii) limited overlap of detection/excitation geometries for labels with large spectral separation. Generally, only few fluorescence-based methods are available to detect ternary or higher order interactions of proteins (15–17). First *in vitro* approaches to perform FCS on more than two species exploited different Stokes shifts of quantum dots (18) or fluorescent dyes excited with a single laser line (19) or two-photon excitation (20, 21), coupled with detection on two or more single photon counting detectors. Following an alternative conceptual approach, it was shown *in vitro* that two spectrally strongly overlapping fluorophore species can be discriminated in FCS by applying statistical filtering of detected photons based on spectrally resolved (fluorescence spectral correlation spectroscopy, FSCS (22)) or fluorescence lifetime (fluorescence lifetime correlation spectroscopy, FLCS (23–25)) detection. Such a framework allows the minimization of cross-talk artefacts in FCCS measurements performed in living cells (26). Recently, three-species implementations of RICCS and FCCS were successfully demonstrated for the first time in living cells. Schrimpf et al. presented raster spectral image correlation spectroscopy (RSICS), a powerful combination of RICS with spectral detection and statistical filtering based on the emission spectra of mEGFP, mVenus and mCherry fluorophores (27). Stefl et al. developed single-color fluorescence lifetime cross-correlation spectroscopy (sc-FLCCS), taking advantage of several GFP variants characterized by short or long fluorescence lifetimes (28). Using this elegant approach, three-species FCCS measurements could be performed in yeast cells, with just two excitation lines.

Here, we explore the full potential of FSCS and RSICS. In particular, we present scanning fluorescence spectral correlation spectroscopy (SFSCS), combining SFCS and FSCS. We show that SFSCS enables cross-talk-free SFCCS measurements of two protein species at the PM of living cells tagged with strongly overlapping fluorophores in the green or red region of the visible spectrum, excited with a single excitation line. This approach results in correct estimates of protein diffusion dynamics, oligomerization, and interactions between both species. Further, we extend our approach to the analysis of three or four interacting partners: by performing cross-correlation measurements on different fluorescent protein (FP) hetero-oligomers, we demonstrate that up to four FP species can be simultaneously analyzed. We then apply this scheme to simultaneously investigate the interaction of influenza A virus (IAV) matrix protein 2 (M2) with two cellular host factors, the tetraspanin CD9 and the autophagosome protein LC3, co-expressed in the same cell. Finally, we extend RSICS for the detection of four molecular species and quantify, for the first time directly in living cells, the complete stoichiometry of ternary IAV polymerase complexes assembling in the nucleus, using three-species fluorescence correlation and brightness analysis.

## RESULTS

### Cross-talk-free scanning SFSCS analysis of membrane-associated proteins using FPs with strongly overlapping emission spectra and a single excitation wavelength

To test the suitability of SFSCS to quantify interactions between membrane proteins tagged with strongly spectrally overlapping fluorophores, we investigated HEK 293T cells co-expressing mp-mEGFP and mp-mEYFP. These monomeric FPs are anchored independently to the inner leaflet of the PM and their emission maxima are only ca. 20 nm apart (Fig.S3). The signal originating from the two fluorophores was decomposed using spectral filters (Fig.S4A) based on the emission spectra detected on cells expressing mp-mEGFP and mp-mEYFP separately (Fig.S3). We then calculated autocorrelation functions (ACFs) and the cross-correlation function (CCF) for signal fluctuations assigned to each fluorophore species. Representative CFs for a typical measurement are shown in Fig.1A, indicating absence of interactions and negligible cross-talk between the two FPs. In contrast, we observed substantial CCFs when analyzing measurements on cells expressing mp-mEYFP-mEGFP hetero-dimers (Fig.S5A). Overall, we obtained a relative cross-correlation (rel.cc.) of 0.72±0.12 (mean±SD, n=22 cells) in the latter sample, compared to a vanishing rel.cc. of 0.02±0.04 (mean±SD, n=34 cells) in the negative control (Fig.1B). Comparison of two types of linker peptides (short flexible or long rigid) between mEGFP and mEYFP showed that the linker length slightly affected rel.cc. values obtained on hetero-dimers (Fig.S5C). FPs linked by a short peptide displayed lower rel.cc., probably due to fluorescence resonance energy transfer (FRET), as previously reported (29). Therefore, unless otherwise noted, similar long rigid linkers were inserted in all constructs used in this study that contain multiple FPs (see Table S2).

**Figure 1.**
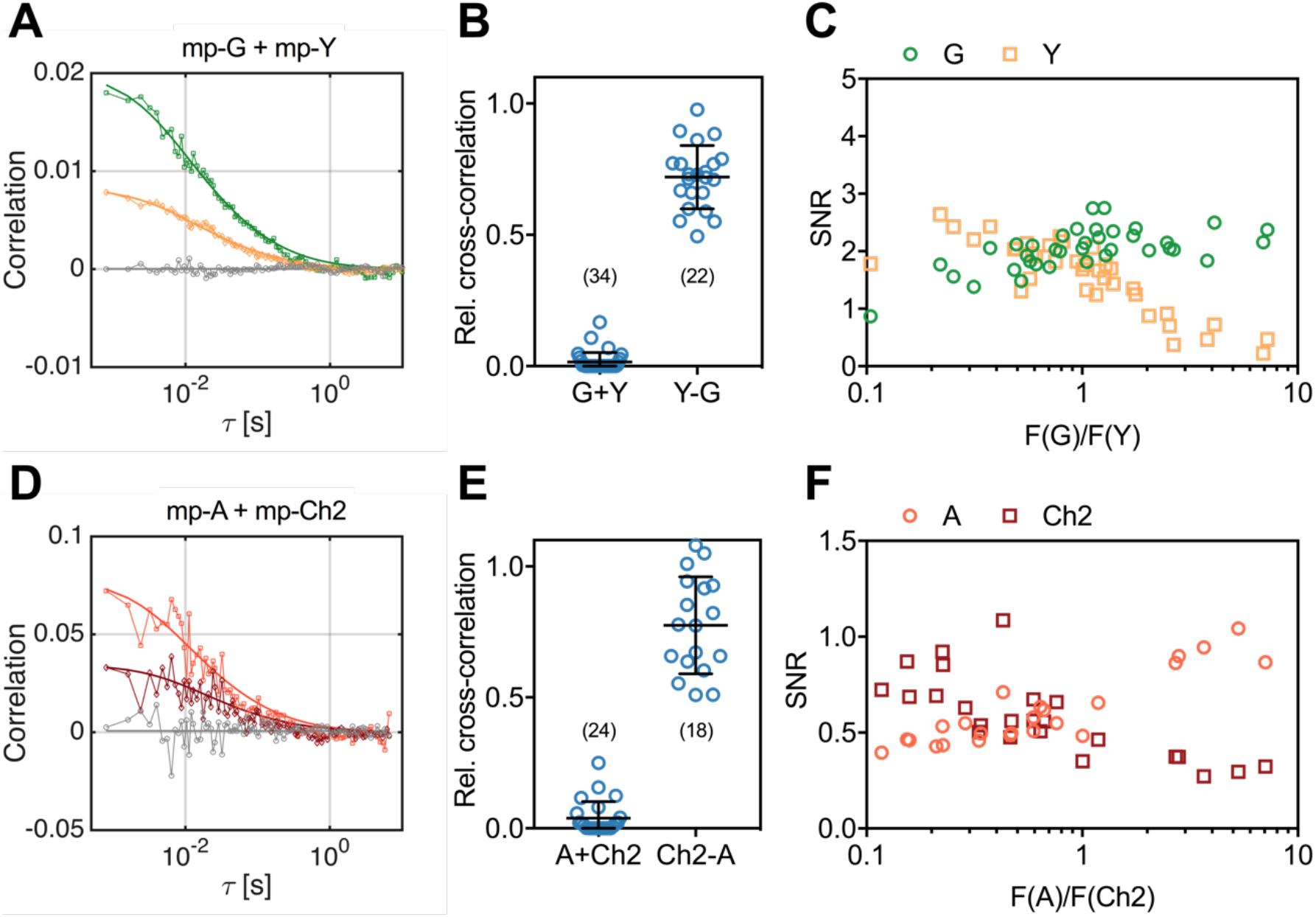
Cross-correlation and SNR analysis for two-species SFSCS measurements at the PM of HEK 293T cells, performed with FPs showing strongly overlapping emission spectra. **(A)** Representative CFs (green: ACF for mEGFP (“G”), yellow: ACF for mEYFP (“Y”), grey: CCF calculated for both fluorophore species) obtained from SFSCS measurements on the PM of HEK 293T cells co-expressing mp-mEGFP and mp-mEYFP. Solid thick lines show fits of a two-dimensional diffusion model to the CFs. **(B)** Relative cross-correlation values obtained from SFSCS measurements described in (A) (“G+Y”) or on HEK 293T cells expressing mp-mEYFP-mEGFP hetero-dimers (“Y-G”). **(C)** SNR of ACFs for mEGFP (green) and mEYFP (yellow), obtained from SFSCS measurements described in (A), plotted as a function of the average ratio of detected mEGFP and mEYFP fluorescence. **(D)** Representative CFs (light red: ACF for mApple (“A”), dark red: ACF for mCherry2 (“Ch2”), grey: CCF calculated for both fluorophores) obtained from SFSCS measurements on the PM of HEK 293T cells co-expressing mp-mApple and mp-mCherry2. Solid thick lines show fits of a two-dimensional diffusion model to the CFs. **(E)** Relative cross-correlation values obtained from SFSCS measurements described in (D) (“A+Ch2”) or on HEK 293T cells expressing mp-mCherry2-mApple hetero-dimers (“Ch2-A”). **(F)** SNR of ACFs for mApple (light red) and mCherry2 (dark red), obtained from SFSCS measurements described in (D), plotted as a function of the average ratio of detected mApple and mCherry2 fluorescence. Data are pooled from three (B) or two (E) independent experiments each. The number of cells measured is given in parentheses. Error bars represent mean±SD.

Overlapping fluorescence emission from different species detected in the same channels provides unwanted background signal and thus reduces the signal-to-noise ratio (SNR) of the CFs (27). To assess to which extent the SNR depends on the relative concentration of mEGFP and mEYFP fluorophores, we compared it between measurements on cells with different relative expression levels of the two membrane constructs (Fig.1C). While the SNR of mEGFP ACFs was only moderately affected by the presence of mEYFP signal (i.e. SNR ranging from ca. 2.5 to 1.0, with 90% to 10% of the signal originating from mEGFP), the ACFs measured for mEYFP showed strong noise when mEGFP was present in much higher amount (i.e. SNR ranging from 2.5 to 0.2, with 90% to 10% of the signal originating from mEYFP).

Next, we tested whether the same approach can be used for FPs with overlapping emission in the red region of the visible spectrum, which generally suffer from reduced SNR in FFS applications (14, 30). Therefore, we performed SFSCS measurements on HEK 293T cells co-expressing mp-mCherry2 and mp-mApple. Also the emission spectra of these FPs are shifted by less than 20 nm (Fig.S3, spectral filters are shown in Fig.S4B). Correlation analysis resulted generally in noisier CFs (Fig.1D) compared to mEGFP and mEYFP. Nevertheless, a consistently negligible rel.cc. of 0.04±0.06 (mean±SD, n=24 cells) was observed. In contrast, a high rel.cc. of 0.78±0.19 (mean±SD, n=18 cells) was obtained on cells expressing mp-mCherry2-mApple hetero-dimers (Fig.1E, Fig.S5B). SNR analysis confirmed lower SNRs of the CFs obtained for red FPs (Fig.1F) compared to mEGFP and mEYFP, with mApple depending more weakly on the relative fluorescence signal than mCherry2 (i.e. ca. 2-fold change for mApple vs. ca. 4-fold change for mCherry2, when the relative abundance changed from 90% to 10%).

We furthermore verified that SFSCS analysis results in correct estimates of protein diffusion dynamics. To this aim, we co-expressed mEGFP-tagged IAV hemagglutinin spike transmembrane protein (HA-mEGFP) and mp-mEYFP. We then compared the diffusion times measured by SFSCS to the values obtained on cells expressing each of the two constructs separately (Fig.2A). For HA-mEGFP, an average diffusion time of 36±8 ms (mean±SD, n=18 cells) was determined in cells expressing both proteins. This value was comparable to that measured for HA-mEGFP expressed separately (34±9 ms, mean±SD, n=21 cells). For mp-mEYFP, diffusion times of 8±2 ms and 9±3 ms were measured in samples expressing both proteins or just mp-mEYFP, respectively. In addition to diffusion analysis, we also analyzed the cross-correlation of HA-mEGFP and mp-mEYFP signal for two-species measurements, resulting in negligible rel.cc. values close to zero (Fig.S6). Hence, SFSCS yielded correct estimates of diffusion dynamics and allowed to distinguish faster and slower diffusing protein species tagged with spectrally strongly overlapping FPs.

**Figure 2.**
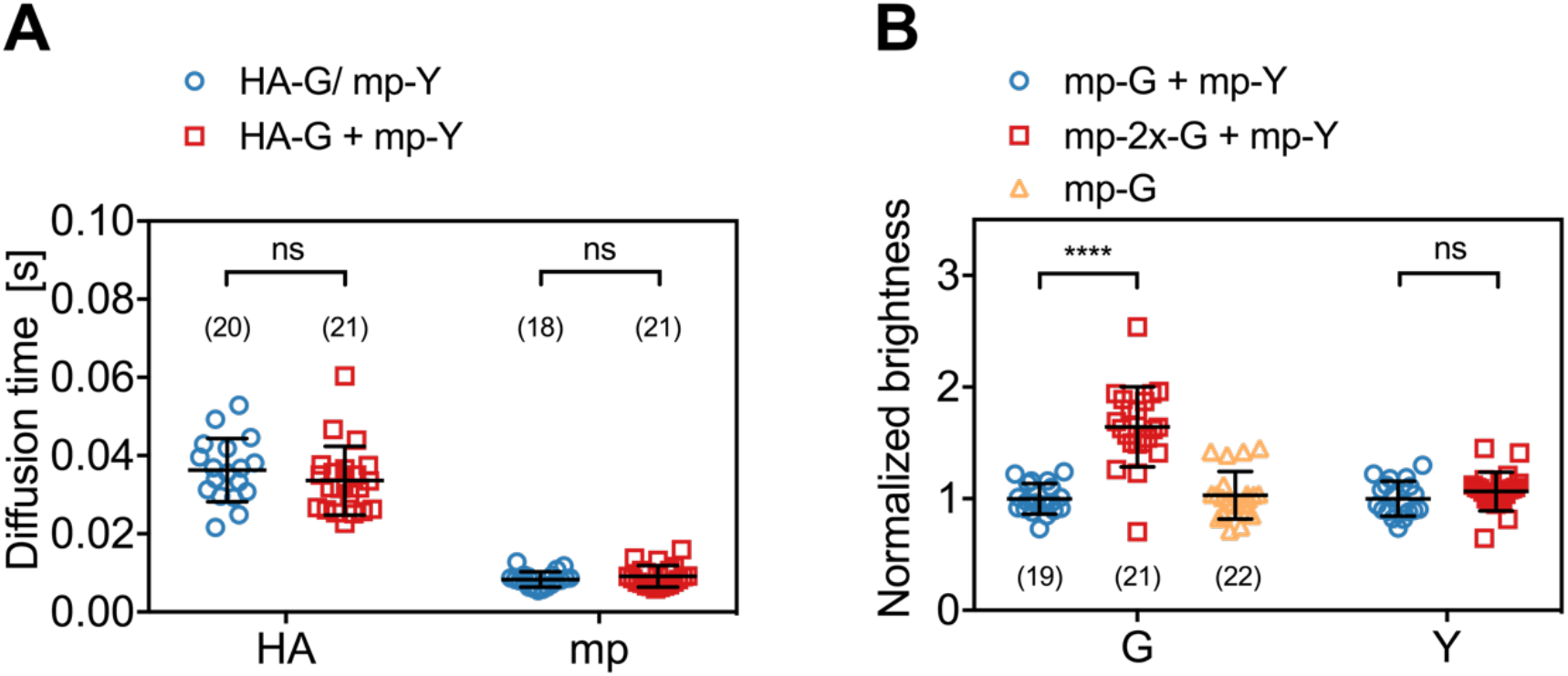
Diffusion and molecular brightness analysis for two-species SFSCS measurements at the PM of HEK 293T cells. **(A)** Diffusion times obtained from SFSCS measurements on HEK 293T cells expressing either IAV HA-mEGFP or mp-mEYFP separately (blue), or co-expressing both fusion proteins (red). **(B)** Normalized molecular brightness values obtained from SFSCS measurements on HEK 293T cells co-expressing mp-mEGFP and mp-mEYFP (blue), mp-2x-mEGFP and mp-mEYFP (red), or expressing mp-mEGFP alone (yellow). Normalized brightness values were calculated by dividing molecular brightness values detected in each SFSCS measurement by the average brightness obtained for mEGFP and mEYFP in cells co-expressing mp-mEGFP and mp-mEYFP. Data are pooled from two independent experiments for each sample. The number of cells measured is given in parentheses. Error bars represent mean±SD. Statistical significance was determined using Welch’s corrected two-tailed student’s *t*-test (\*\*\*\**P*<0.0001, ns: not significant).

Finally, we evaluated the capability of SFSCS to precisely determine the molecular brightness, as a measure of protein oligomerization. We compared the molecular brightness values for mEGFP and mEYFP in samples co-expressing monomeric FP constructs mp-mEGFP and mp-mEYFP with the values obtained for cells co-expressing mp-2x-mEGFP homo-dimers and mp-mEYFP (Fig.2B). From SFSCS analysis of measurements in the latter sample, we obtained a normalized molecular brightness of 1.64±0.36 (mean±SD, n=21 cells) for mEGFP, relative to the brightness determined in the monomer sample (n=19 cells). This value is in agreement with our previous quantification of the relative brightness of mEGFP homo-dimers, corresponding to a fluorescence probability (*p*_*f*_) of ca. 60-75% for mEGFP (14). The *p*_*f*_ is an empirical, FP-specific parameter that was previously characterized for multiple FPs (14). It quantifies the fraction of non-fluorescent FPs due to photophysical processes, such as transitions to long-lived dark states, or slow FP maturation and needs to be taken into account to correctly determine the oligomerization state of FP tagged protein complexes. As a reference for the absolute brightness, we also determined the relative molecular brightness of mEGFP in cells expressing mp-mEGFP alone, yielding a value of 1.03±0.21 (mean±SD, n=22 cells). Additionally, the brightness values determined for mEYFP in both two-species samples were similar, with a relative ratio of 1.07±0.18, as expected. This confirms that reliable brightness values were obtained and that dimeric and monomeric species can be correctly identified.

In summary, these results demonstrate that SFSCS analysis of fluorescence fluctuations successfully separates the contributions of FPs exhibiting strongly overlapping emission spectra, yielding correct quantitative estimates of protein oligomerization and diffusion dynamics.

### Simultaneous cross-correlation and brightness analysis for three spectrally overlapping FPs at the PM

In the previous section, we showed that SFSCS enables cross-talk-free cross-correlation analysis of two fluorescent species excited with a single laser line, even in the case of strongly overlapping emission spectra. To explore the full potential of SFSCS, we extended the approach to systems containing three spectrally overlapping fluorophores. We excited mEGFP, mEYFP, and mCherry2 with 488 nm and 561 nm lines simultaneously and detected their fluorescence in 23 spectral bins in the range of 491 nm to 695 nm. We measured individual emission spectra (Fig.S3) for single species samples to calculate three-species spectral filters (Fig.S4C), which we then used to decompose the signal detected in cells expressing multiple FPs into the contribution of each species.

As a first step, we performed three-species SFSCS measurements on HEK 293T cells co-expressing mp-mEYFP with either i) mp-mEGFP and mp-mCherry2 (mp-G + mp-Y + mp-Ch2) or ii) mp-mCherry2-mEGFP hetero-dimers (mp-Ch2-G + mp-Y). Additionally, we tested a sample with cells expressing mp-mEYFP-mCherry2-mEGFP hetero-trimers (mp-Y-G-Ch2). We then calculated ACFs for all three FP species and CCFs all fluorophore combinations, respectively. In the first sample (mp-G + mp-Y + mp-Ch2), in which all three FPs are anchored independently to the PM, we obtained CCFs fluctuating around zero for all fluorophore combinations, as expected (Fig.3A). In the second sample (mp-Ch2-G + mp-Y), a substantial cross-correlation was detected between mEGFP and mCherry2, whereas the other two combinations resulted in CCFs fluctuating around zero (Fig.3B). In the hetero-trimer sample, CCFs with low level of noise and amplitudes significantly above zero were successfully obtained for all three fluorophore combinations (Fig.3C). From the amplitude ratios of the ACFs and CCFs, we then calculated rel.cc. values for all measurements (Fig.3F). Negligible rel.cc. values were obtained for all fluorophore combinations that were not expected to show interactions, e.g. 0.05±0.08 (mean±SD, n=46 cells) between mEGFP and mEYFP signal in the first sample. Low residual rel.cc. values obtained in a few measurements are likely an artefact of the fitting procedure (caused by restriction of the correlation amplitude to positive values). For noisy CCFs, this issue is responsible for low, but positive correlation amplitudes (and thus rel.cc. values) even in the absence of significant correlation (i.e. for CCFs fluctuating around zero), and a high uncertainty of this fit parameter. For mEGFP and mCherry2, similar rel.cc. values of 0.45±0.06 (mean±SD, n=20 cells) and 0.56±0.08 (mean±SD, n=17 cells) were observed in cells expressing mp-mCherry2-mEGFP hetero-dimers or mp-mEYFP-mCherry2-mEGFP hetero-trimers. The minor difference could be attributed to different linker peptides (i.e. long rigid linker between FPs in hetero-trimers and a short flexible linker in hetero-dimers), increasing the degree of FRET between mEGFP and mCherry2 in hetero-dimers and reducing the cross-correlation. The hetero-trimer sample showed high rel.cc. values also for the other two fluorophore combinations: mEGFP and mEYFP (rel.cc._G,Y_=0.79±0.12) or mCherry2 and mEYFP (rel.cc._Y,Ch2_=0.57±0.07).

**Figure 3.**
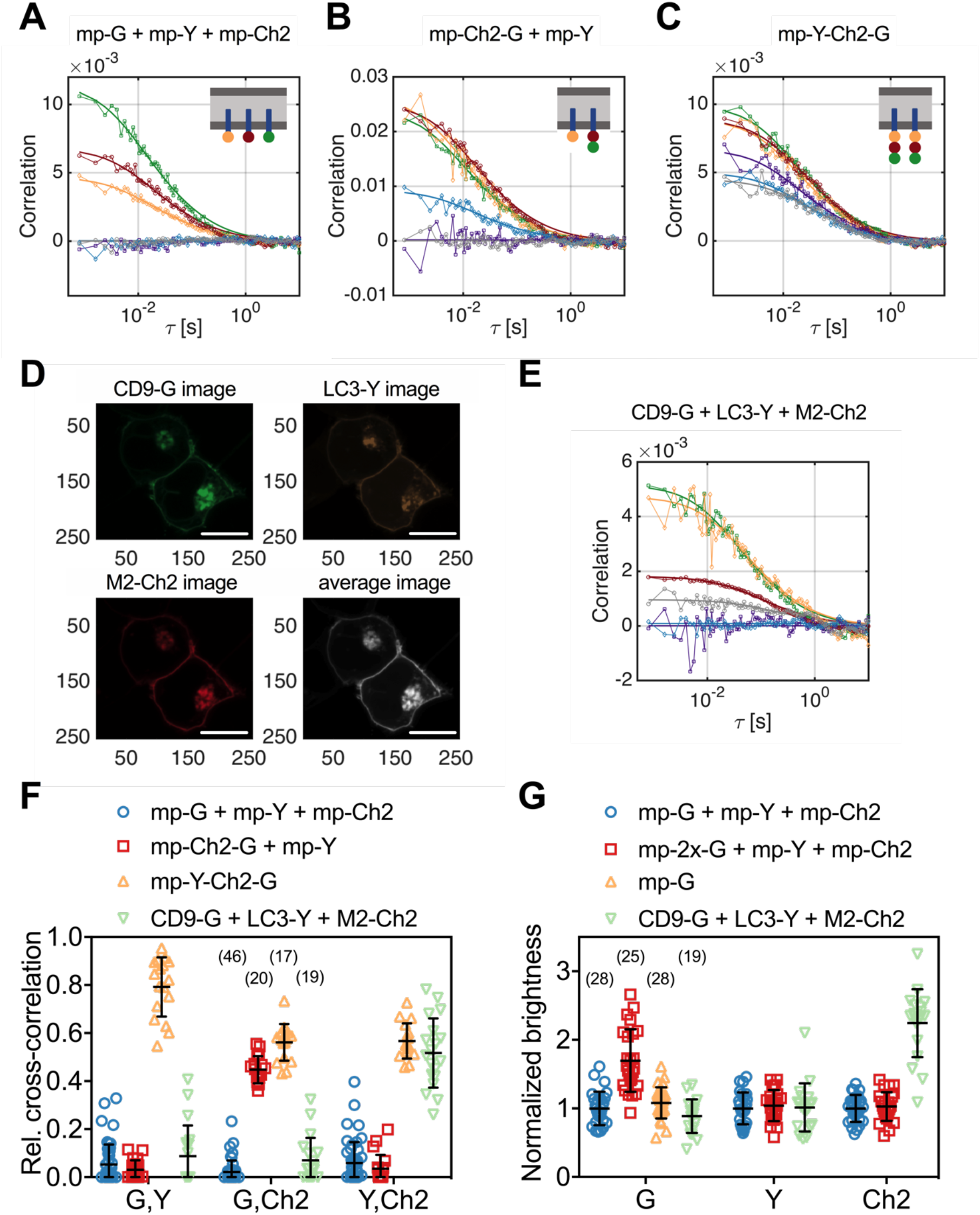
Cross-correlation and molecular brightness analysis for three-species SFSCS measurements on FP hetero-oligomers and IAV M2 at the PM of HEK 293T cells. **(A-C)** Representative CFs (green/yellow/red: ACFs for mEGFP (“G”)/ mEYFP (“Y”)/ mCherry2 (“Ch2”), purple/blue/grey: CCFs calculated for the pairs mEGFP and mEYFP/ mEGFP and mCherry2/ mEYFP and mCherry2) obtained from three-species SFSCS measurements on HEK 293T cells co-expressing mp-mEGFP, mp-mEYFP, and mCherry2 (A), mp-mCherry2-mEGFP hetero-dimers and mp-mEYFP (B), or expressing mp-mEYFP-mCherry2-mEGFP hetero-trimers (C), as illustrated in insets. Solid thick lines show fits of a two-dimensional diffusion model to the CFs. **(D)** Representative fluorescence images of HEK 293T cells co-expressing CD9-mEGFP, LC3-mEYFP, and IAV protein M2-mCh2. Spectral filtering and decomposition were performed to obtain a single image for each species. Scale bars are 5 μm. **(E)** Representative CFs (green/yellow/red: ACFs for mEGFP/mEYFP/mCherry2, purple/blue/grey: CCFs calculated for the pairs mEGFP and mEYFP/ mEGFP and mCherry2/ mEYFP and mCherry2) obtained from three-species SFSCS measurements on HEK 293T cells co-expressing CD9-mEGFP, LC3-mEYFP, and M2-mCh2. Solid thick lines show fits of a two-dimensional diffusion model to the CFs. **(F)** Relative cross-correlation values obtained from three-species SFSCS measurements described in (A-C) and I. **(G)** Normalized molecular brightness values obtained from three-species SFSCS measurements on HEK 293T cells co-expressing mp-mEGFP, mp-mEYFP, and mp-mCherry2 (blue), mp-2x-mEGFP, mp-mEYFP, and mp-mCherry2 (red), CD9-mEGFP, LC3-mEYFP, and M2-mCh2 (green), or expressing mp-mEGFP alone (yellow). Normalized brightness values were calculated by dividing the molecular brightness values detected in each SFSCS measurement by the average brightness obtained for mEGFP, mEYFP, and mCherry2 in cells co-expressing mp-mEGFP, mp-mEYFP, and mp-mCherry2. Data are pooled from two independent experiments for each sample. The number of cells measured is given in parentheses. Error bars represent mean±SD.

In addition to cross-correlation analysis, we performed molecular brightness measurements on samples containing three FP species. In particular, we compared molecular brightness values obtained by SFSCS on HEK 293T cells co-expressing homo-dimeric mp-2x-mEGFP, mp-mEYFP, and mp-mCherry2 (mp-2x-G + mp-Y + mp-Ch2) to the values measured on cells co-expressing the three monomeric constructs mp-mEGFP, mp-mEYFP, and mp-mCherry2 (mp-G + mp-Y + mp-Ch2). Whereas similar brightness values were obtained for mEYFP and mCherry2 in both samples, e.g. relative brightness of 1.04±0.23 for mEYFP and 1.03±0.21 for mCherry2 (mean±SD, n=25 cells/ n=28 cells), a higher brightness of 1.70±0.46 was measured for mEGFP in the first sample (Fig.3G). This value corresponds to a *p*_*f*_ of ca. 70% for mEGFP, as expected (14). To confirm that absolute brightness values are not influenced by the spectral decomposition, we also determined the brightness of mEGFP in cells expressing mp-mEGFP alone (Fig.3G), resulting in values close to 1 (1.08±0.23, mean±SD, n=28 cells).

### The IAV protein M2 interacts strongly with LC3 but not with CD9

Having demonstrated the capability of SFSCS to successfully quantify protein interactions and oligomerization, even in the case of three FPs with overlapping emission spectra, we applied this approach in a biologically relevant context. In more detail, we investigated the interaction of IAV channel protein M2 with the cellular host factors CD9 and LC3. CD9 belongs to the family of tetraspanins and is supposedly involved in virus entry and virion assembly (31–33). The autophagy marker protein LC3 was recently shown to be recruited to the PM in IAV-infected cells (see also Fig.S7A,B), promoting filamentous budding and virion stability, thus indicating a role of LC3 in virus assembly (34). To detect hetero-interactions between CD9, LC3 and M2, we co-expressed the fluorescent fusion proteins CD9-mEGFP, LC3-mEYFP and M2-mCherry2 (i.e. M2 carrying an mCherry2 tag at the extracellular terminus) in HEK 293T cells (Fig.3D) and performed three-species SFSCS measurements at the PM (Fig.3E).

We then calculated rel.cc. values to quantify pair-wise interactions of the three proteins (Fig.3F). The obtained rel.cc. values for CD9-mEGFP with LC3-mEYFP or M2-mCherry2 (rel.cc._CD9-G,LC3-Y_=0.09±0.13,rel.cc._CD9-G,M2-Ch2_=0.07±0.09, mean±SD, n=19 cells) were similar to those of the negative cross-correlation control (i.e. cells co-expressing mp-mEGFP, mp-mEYFP and mp-mCherry2, see previous paragraph). In contrast, we detected a substantial rel.cc. of 0.52±0.14 for LC3-mEYFP and M2-mCherry2. This value was close (ca. 90% on average) to that obtained for this fluorophore combination in measurements on FP hetero-trimers, suggesting very strong association of LC3-mEYFP with M2-mCherry2. We furthermore analyzed the molecular brightness for each species, normalized to the monomeric references (Fig.3G). While CD9-mEGFP and LC3-mEYFP showed normalized brightness values close to 1 (B_CD9-G_=0.89±0.25, B_LC3-Y_=1.02±0.35), suggesting that both proteins are monomers, we observed significantly higher relative brightness values for M2-mCherry2 (B_M2-Ch2_=2.24±0.49). Assuming a *p*_*f*_ of ca. 60% for mCherry2 (14), the determined relative brightness corresponds to an oligomerization state of *ε*_*M2-Ch2*_ = 3.1 ± 0.8, i.e. formation of M2 dimers to tetramers at the PM.

### SFSCS allows simultaneous analysis of protein-protein interactions for four spectrally overlapping FP species

Having demonstrated robust three-species cross-correlation analysis, we aimed to further explore the limits of SFSCS. We investigated therefore whether SFSCS can discriminate of differential interactions between four species using the spectral emission patterns of mEGFP, mEYFP, mApple and mCherry2 for spectral decomposition (Fig.S3, Fig.S4D). As a proof of concept, we performed four-species measurements on three different samples: i) cells co-expressing all four FPs independently as membrane-anchored proteins (mp-G + mp-Y + mp-A + mp-Ch2), ii) cells co-expressing mp-mCherry2-mEGFP hetero-dimers, mp-mEYFP, and mp-mApple (mp-Ch2-G + mp-Y + mp-A), iii) cells expressing mp-mEYFP-mCherry2-mEGFP-mApple hetero-tetramers (mp-Y-Ch2-G-A). We then calculated four ACFs, six CCFs, and rel.cc. values from the amplitude ratios of the ACFs and CCFs. For all fluorophore species, ACFs with amplitudes significantly above zero were obtained. ACFs calculated for mEGFP and mEYFP were characterized by a higher SNR compared to those for the red FPs mApple and, in particular, mCherry2 (Fig.4A-C). Nevertheless, reasonable diffusion time values could be determined for all species, showing the largest variation for mCherry2 (Fig.S8).

**Figure 4.**
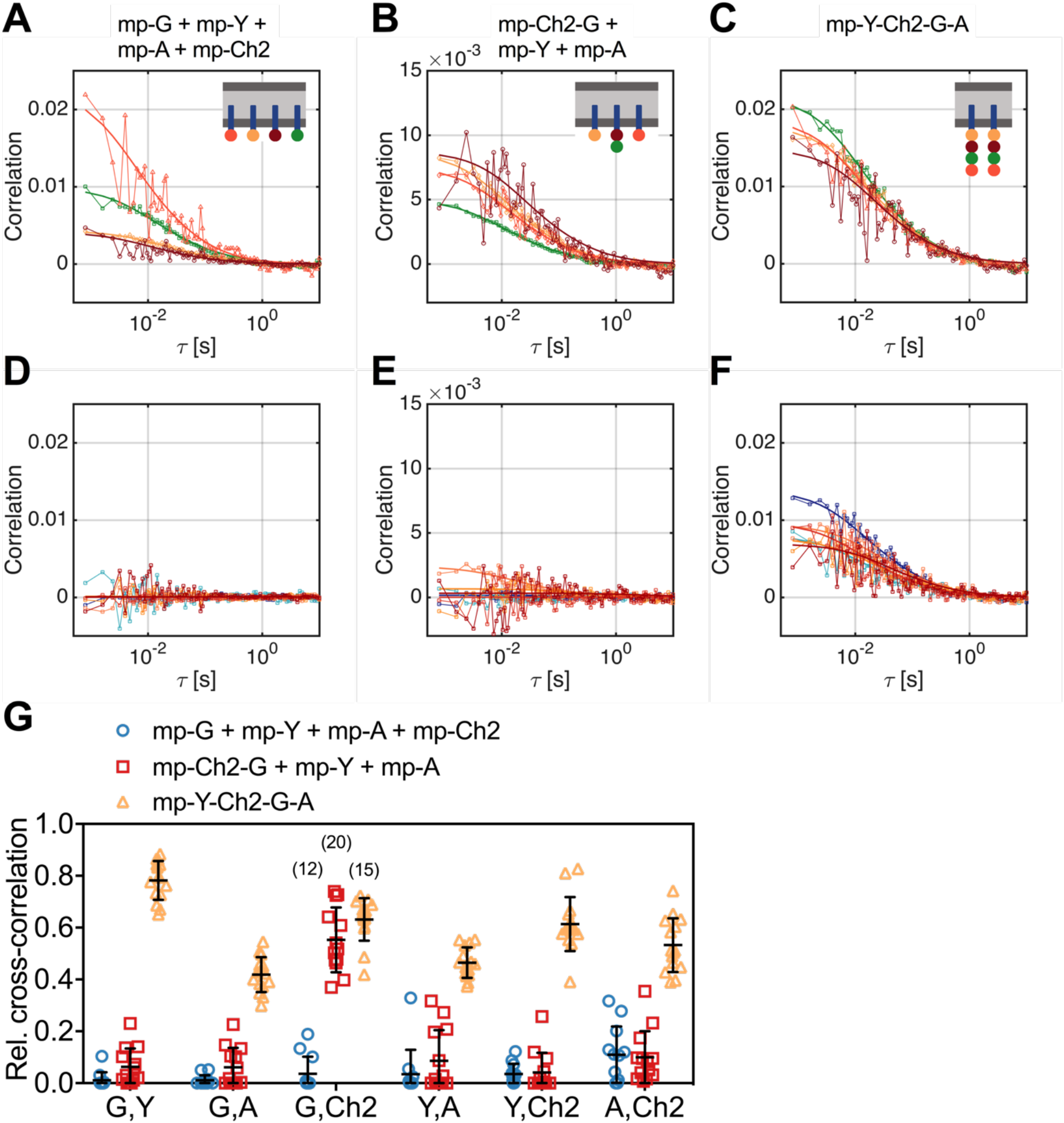
Cross-correlation analysis for four-species SFSCS measurements on FP hetero-oligomers in HEK 293T cells. **(A-C)** Representative ACFs (green/yellow/orange/red for mEGFP (“G”)/ mEYFP (“Y”)/ mApple (“A”)/ mCherry2 (“Ch2)) obtained from four-species SFSCS measurements on HEK 293T cells co-expressing mp-mEGFP, mp-mEYFP, mp-mApple, and mCherry2 (A), mp-mCherry2-mEGFP hetero-dimers, mp-mEYFP, and mp-mApple (B), or expressing mp-mEYFP-mCherry2-mEGFP-mApple hetero-tetramers (C), as illustrated in insets. Solid thick lines show fits of a two-dimensional diffusion model to the CFs. **(D-F)** SFSCS CCFs (dark blue/ light blue/ orange/ yellow/ red/ dark red for CCFs calculated for mEGFP and mEYFP/ mEGFP and mApple/ mEGFP and mCherry2/ mEYFP and mApple/ mEYFP and mCherry2/ mApple and mCherry2) from measurements described in (A-C) (CCFs in (I(E)/(F) corresponding to ACFs shown in (A)/(B)/(C)). Solid thick lines show fits of a two-dimensional diffusion model to the CFs. **(G)** Relative cross-correlation values obtained from four-species SFSCS measurements described in (A-C). Data are pooled from two independent experiments. The number of cells measured is given in parentheses. Error bars represent mean±SD.

Noise levels of the CCFs were moderate (Fig.4D-F), yet allowing robust fitting and estimation of cross-correlation amplitudes. Based on the determined rel.cc. values (Fig.4G), the different samples could successfully be discriminated. In the first sample (mp-G + mp-Y + mp-A + mp-Ch2), neglibile to very low values were obtained, i.e. at maximum 0.11±0.11 (mean±SD, n=12 cells) for mApple and mCherry2. In the second sample (mp-Ch2-G + mp-Y + mp-A), similarly low rel.cc. values were obtained for all fluorophore combinations, e.g. 0.10±0.10 (mean±SD, n=13 cells) for mApple and mCherry2, with the exception of mEGFP and mCherry2, showing an average value of 0.55±0.13. For the hetero-tetramer sample, high rel.cc. values were measured for all fluorophore combinations, ranging from 0.42±0.07 (mean±SD, n=15 cells) for mEGFP and mApple to 0.78±0.08 for mEGFP and mEYFP. Notably, a significant rel.cc. of 0.53±0.10 was also determined for mApple and mCherry2 signals, i.e. from the CCFs exhibiting the lowest SNR.

### RSICS can be extended to simultaneous detection of four fluorophore species

Having identified a set of FPs that is compatible with four-species SFSCS, we aimed to extend the recently presented RSICS method (27) to applications with four fluorophore species being detected simultaneously. To test the effectiveness of this approach, we carried out measurements in living A549 cells co-expressing mEGFP, mEYFP, mApple, and mCherry2 in several configurations, similar to the SFSCS experiments presented in the previous paragraph. In more detail, we performed four-species RSICS measurements on the following three samples: i) cells co-expressing free mEGFP, mEYFP, mApple, and mCherry2 (1x-G + 1x-Y + 1x-A + 1x-Ch2), ii) cells co-expressing mCherry2-mEGFP and mEYFP-mApple hetero-dimers (Ch2-G + Y-A), iii) cells expressing mEYFP-mCherry2-mEGFP-mApple hetero-tetramers (Y-Ch2-G-A). Representative CFs obtained following RSICS analysis with arbitrary region selection (35) are shown in Fig.5. In all samples, ACFs with amplitudes significantly above zero were obtained, with the highest noise level detected for mCherry2 (Fig.5A,C,E). A three-dimensional diffusion model could be successfully fitted to all detected ACFs.

**Figure 5.**
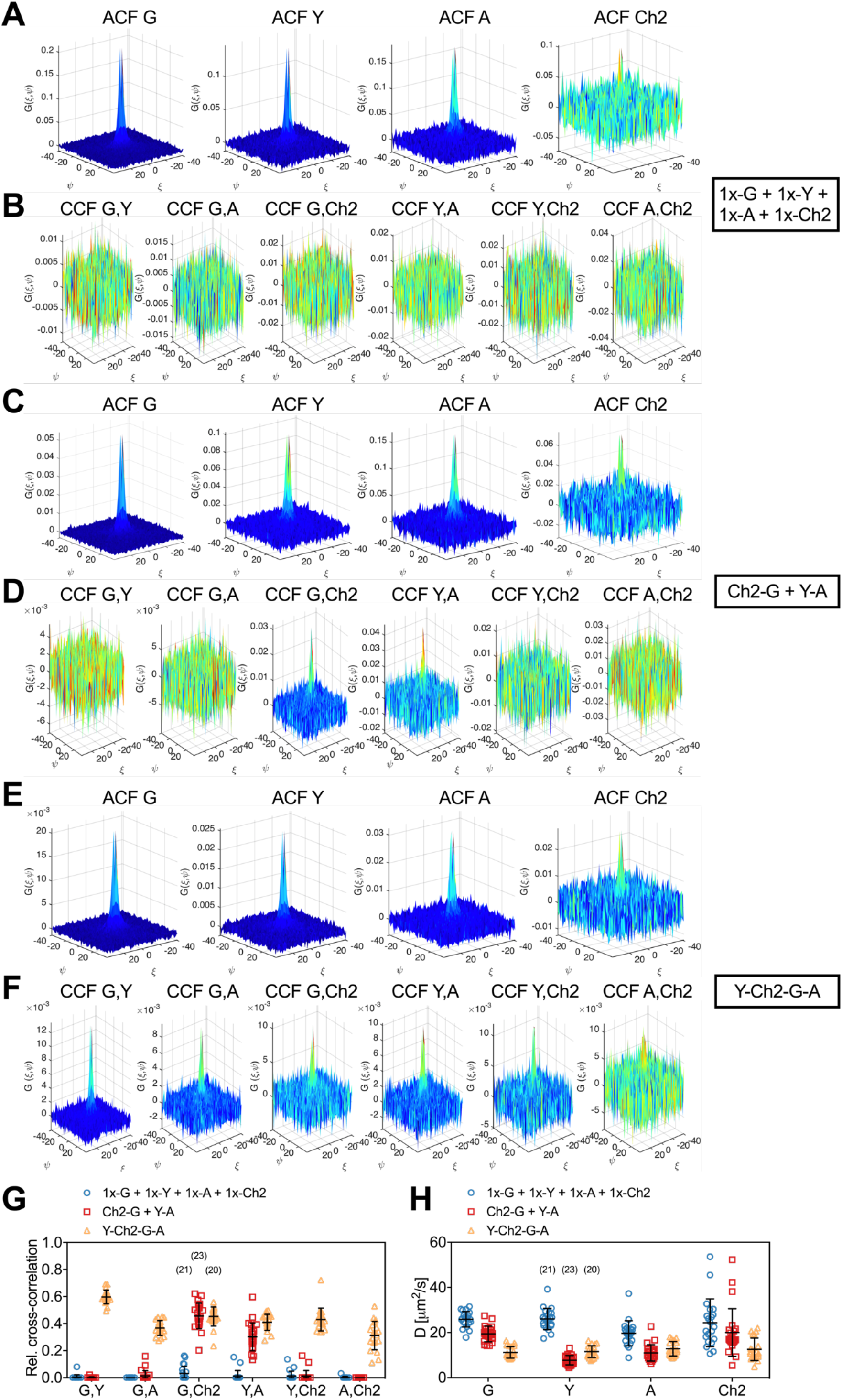
Cross-correlation analysis for four-species RSICS measurements on FP hetero-oligomers expressed in A549 cells. **(A-F)** Representative RSICS spatial ACFs (A,C,E) and CCFs (B,D,F) obtained from four-species RSICS measurements on A549 cells. Cells were co-expressing mEGFP (“G”), mEYFP (“Y”), mApple (“A”), mCherry2 (“Ch2”) (A,B), mCherry2-mEGFP and mEYFP-mApple heterodimers (C,D), or mEYFP-mCherry2-mEGFP-mApple hetero-tetramers (E,F). **(G,H)** Relative cross-correlation values (G) and diffusion coefficients (H) obtained from four-species RSICS measurements described in (A-F). Data are pooled from two independent experiments. The number of cells measured is given in parentheses. Error bars represent mean±SD.

Detected CCFs showed the expected pattern: all six CCFs were indistinguishable from noise for the first sample with four independent FPs (Fig.5B), whereas large CCF amplitudes were obtained for the pairs mEGFP and mCherry2, as well as mEYFP and mApple in the second sample (Ch2-G + Y-A) (Fig.5D). Also, significantly large amplitudes were observed for all six CCFs for the hetero-tetramer sample, albeit with different levels of noise. For example, the lowest SNR was observed in CCFs for mApple and mCherry2 (Fig.5F).

From the amplitude ratios of ACFs and CCFs, we determined rel.cc. values (Fig.5G). This analysis resulted in negligible values for the first sample (1x-G + 1x-Y + 1x-A + 1x-Ch2), e.g. rel.cc._G,Ch2_=0.03±0.05 (mean±SD, n=21 cells). For the second sample (Ch2-G + Y-A), values significantly above zero, i.e. rel.cc._G,Ch2_=0.46±0.09 (mean±SD, n=23 cells) and rel.cc._Y,A_=0.30±0.10, were only observed for two fluorophore pairs. For the third sample, cells expressing mEYFP-mCherry2-mEGFP-mApple hetero-tetramers (Y-Ch2-G-A), rel.cc. values significantly above zero were obtained for all FP pairs, ranging from rel.cc._A,Ch2_=0.31±0.11 (mean±SD, n=20 cells) to rel.cc._G,Y_=0.60±0.05. Notably, rel.cc. values obtained for the FP species correlating in the second sample (Ch2-G + Y-A) were similar in the third sample (Y-Ch2-G-A), e.g. rel.cc._G,Ch2_=0.45±0.07 and rel.cc._Y,A_=0.41±0.06. The lower rel.cc. value measured for mEYFP and mApple in hetero-dimers (Ch2-G + Y-A) could be attributed to different linker sequences (long rigid linker in hetero-dimers vs. mCherry2-mEGFP and three long rigid linkers as spacer in hetero-tetramers (Y-Ch2-G-A)), possibly affecting FRET between neighboring FPs.

Finally, we analyzed the diffusion dynamics of FP fusion proteins as determined from the spatial dependence of the ACFs for the four fluorophore species. Diffusion coefficients (D) obtained for mCherry2 showed the highest variation (Fig.5H), reflecting the reduced SNR for this fluorophore. Nevertheless, similar average D values were determined for different fluorophore species coupled as hetero-oligomers, e.g. D_G_=19.4±3.4 μm^2^/s and D_Ch2_=20±11 μm^2^/s (mean±SD, n=23 cells) for mEGFP-mCherry2 hetero-dimers, and D_G_=11.2±2.5 μm^2^/s, D_Y_=11.6±2.6 μm^2^/s, D_A_=12.8±3.2 μm^2^/s, D_Ch2_=12.6±5.0 μm^2^/s (mean±SD, n=20 cells) for hetero-tetramers.

### Cross-correlation and molecular brightness analysis via three-species RSICS provide stoichiometry of IAV polymerase complex assembly

To test the versatility of three-species RSICS, we quantified intracellular protein interactions and stoichiometries in a biologically relevant context. As an example, we focused on the assembly of the IAV polymerase complex (PC), consisting of the three subunits polymerase acidic protein (PA), polymerase basic protein 1 (PB1), and 2 (PB2). A previous investigation using FCCS suggested an assembly model in which PA and PB1 form hetero-dimers in the cytoplasm of cells. These are imported into the nucleus and appear to interact with PB2 to form hetero-trimeric complexes (36). Nevertheless, this analysis could only be performed between two of the three subunits at the same time. Also, the stoichiometry of the complex was reported only for one of the three subunits, i.e. PA protein dimerization. Here, we labeled all three subunits using FP fusion constructs and co-expressed PA-mEYFP, PB1-mEGFP, and PB2-mCherry2 in A549 cells. We then performed three-species RSICS measurements in the cell nucleus, where all three proteins are enriched (Fig.6A). RSICS analysis was performed on an arbitrarily-shaped homogeneous region of interest in the nucleus. We then calculated RSICS ACFs (Fig.6B), CCFs (Fig.6C), and rel.cc. values (Fig.6D) for the three fluorophore combinations. The determined rel.cc. values were compared to the values obtained on negative controls (i.e. cells co-expressing free mEGFP, mEYFP, and mCherry) and positive controls (i.e. cells expressing mEYFP-mCherry2-mEGFP hetero-trimers) (Fig.6D).

**Figure 6.**
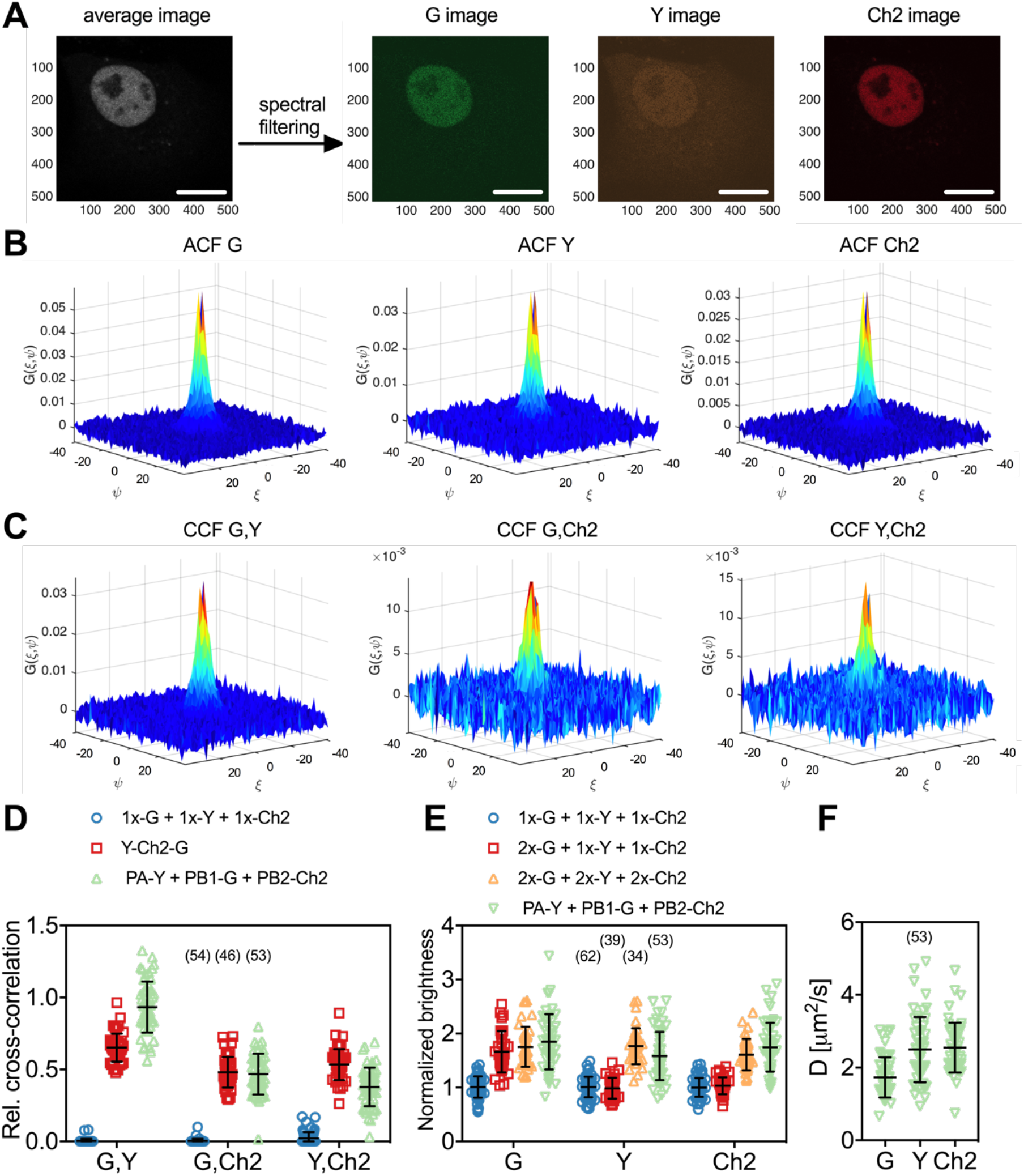
Three-species RSICS measurements on IAV polymerase complex and FP hetero-oligomers in the nucleus of A549 cells. **(A)** Representative fluorescence image (left) of A549 cells co-expressing FP-tagged IAV PC proteins PA-mEYFP, PB1-mEGFP, and PB2-mCherry2. Spectral filtering and decomposition result in a single image for each species (right), denoted with “Y”, “G”, and “Ch2”. Scale bars are 10 μm. **(B,C)** Representative RSICS spatial ACFs (B) and CCFs (C) obtained from three-species RSICS measurements on A549 cells co-expressing PA-mEYFP, PB1-mEGFP, and PB2-mCherry2. **(D)** Relative cross-correlation values obtained from three-species RSICS measurements on A549 cells co-expressing mEGFP, mEYFP, and mCherry2 (blue), PA-mEYFP, PB1-mEGFP, PB2-mCherry2 (green), or expressing mEYFP-mCherry2-mEGFP hetero-trimers (red). Data are pooled from four independent experiments. **(E)** Normalized molecular brightness values obtained from three-species RSICS measurements on A549 cells co-expressing mEGFP, mEYFP, and mCherry2 (blue), 2x-mEGFP, mEYFP, and mCherry2 (red), 2x-mEGFP, 2x-mEYFP, 2x-mCherry2 (yellow), or PA-mEYFP, PB1-mEGFP, and PB2-mCherry2 (green). Data are pooled from three (2x-mEGFP + mEYFP + mCherry2, 2x-mEGFP + 2x-mEYFP + 2x-mCherry2), four (PA-mEYFP + PB1-mEGFP + PB2-mCherry2), or five (mEGFP + mEYFP + mCherry2) independent experiments. **(F)**Diffusion coefficients obtained from three-species RSICS measurements on A549 cells co-expressing PA-mEYFP, PB1-mEGFP, and PB2-mCherry2. Data are pooled from four independent experiments. For (D)-(F), the number of cells measured is given in parentheses. Error bars represent mean±SD.

For the polymerase sample, high rel.cc. values were observed for all combinations: rel.cc._PB1-G,PA-Y_=0.93±0.18 (mean±SD, n=53 cells), rel.cc._PB1-G,PB2-Ch2_=0.47±0.14, rel.cc._PA-Y,PB2-Ch2_=0.39±0.14. For the positive control, similar values were observed for mEGFP and mCherry2, rel.cc._G,Ch2_=0.48±0.11 (mean±SD, n=46 cells), whereas the values were higher than that measured for PCs for mEYFP and mCherry2, rel.cc._Y,Ch2_=0.53±0.11, and lower for mEGFP and mEYFP, rel.cc._G,Y_=0.65±0.10. The lower average rel.cc. between PA-mEYFP and PB2-mCherry2 compared to the positive control indicates the presence of a minor fraction of non-interacting PA and PB2. These proteins could be present in the nucleus in unbound form when expressed in higher amount than PB1, since both PA and PB2 localize in the nucleus individually and were previously shown not to interact when both present without PB1 (36). This explanation is supported by the correlation between rel.cc._PA-Y,PB2-Ch2_ and the relative abundance of PB1-mEGFP (Fig.S9A). Also, the observation that PB1 is only transported to the nucleus in complex with PA is confirmed by the lower concentration of PB1-mEGFP compared to PA-mEYFP in the nuclei of all measured cells (Fig.S9A). Thus, the fraction of PB1-mEGFP bound to PA-mEYFP should be as high as the positive control, for a 1:1 stoichiometry. The observation of higher rel.cc. between mEGFP and mEYFP for the polymerase subunits indicates higher order interactions, i.e. higher stoichiometry than 1:1 (37).

To quantify the stoichiometry of the PC directly, we analyzed the molecular brightness of RSICS measurements for all three fluorophore species. We normalized the obtained values to the average values determined by RSICS on cells co-expressing monomeric mEGFP, mEYFP, and mCherry2, measured on the same day. To test whether RSICS can be used to obtain reliable brightness/oligomerization values for all fluorophore species, we first performed control experiments on cells co-expressing either i) 2x-mEGFP homo-dimers with mEYFP and mCherry monomers (2x-G + 1x-Y + 1x-Ch2) or ii) the three homo-dimers 2x-mEGFP, 2x-mEYFP, and 2x-mCherry2 (2x-G + 2x-Y + 2x-Ch2). In the first sample, we observed an increased relative brightness of 1.67±0.38 (mean±SD, n=34 cells) for mEGFP, whereas values around 1 were obtained for mEYFP and mCherry2. This confirmed the presence of mEGFP dimers as well as mEYFP and mCherry2 monomers in this control sample, as expected (Fig.6E). In the sample containing all three homo-dimers, increased relative brightness values were observed for all fluorophore species: 1.75±0.37 (mean±SD, n=39 cells) for mEGFP, 1.77±0.33 for mEYFP, and 1.61±0.29 for mCherry2. These values indicate successful determination of the dimeric state of all three FP homo-dimers and are in good agreement with previous brightness measurements on homo-dimers of mEGFP, mEYFP and mCherry2, corresponding to *p*_*f*_ values of 60-75% (14). Next, we proceeded with the analysis of PC oligomerization. For each polymerase subunit, relative brightness values close to the values of homo-dimers were observed. Assuming *p*_*f*_ values of 75%, 77%, and 61% (as calculated from the determined relative brightness values of homo-dimers) for mEGFP, mEYFP, and mCherry2, respectively, *p*_*f*_ corrected normalized brightness values of *ε*_*PB1-G*_ = 2.1 ± 0.7 (mean±SD, n=53 cells), *ε*_*PA-Y*_ = 1.8 ± 0.6, and *ε*_*PB2-Ch2*_ = 2.2 ± 0.7 were obtained (see methods for details). These results suggest a 2:2:2 stoichiometry of the IAV PC subunits. Finally, we analyzed the diffusion dynamics of PCs via RSICS (Fig.6F). The average D measured for PB1-mEGFP, D_PB1-G_=1.7±0.6 μm^2^/s (mean±SD, n=53 cells), was ca. 30% lower than the diffusion coefficients determined for PA-mEYFP- and PB2-mCherry2 (D_PA-Y_=2.5±0.9 μm^2^/s and D_PB2-Ch2_=2.6±0.7 μm2/s). This observation is compatible with the above-mentioned presence of a minor fraction of unbound (and thus faster diffusing) PA and PB2 (likely in cells with a lower amount of PB1). A more detailed analysis of the data confirmed this interpretation: The molecular brightness and diffusion coefficient of PA-mEYFP depended on the relative concentration of PB1-mEGFP and PA-mEYFP, i.e. lower brightness and higher diffusion coefficients were obtained in cells where PA-mEYFP was present at much higher concentrations than PB1-mEGFP (Fig.S7B,C).

### Triple raster image correlation spectroscopy (TRICS) analysis provides direct evidence for assembly of ternary IAV polymerase complexes

To directly confirm that IAV PC subunits form ternary complexes in the cell nucleus, we implemented a triple-correlation analysis (i.e. TRICS) to detect coincident fluctuations of the signal emitted by mEGFP-, mEYFP- and mCherry2-tagged proteins. A similar analysis has previously been presented for three-channel FCS measurements (e.g. fluorescence triple correlation spectroscopy (21), triple-color coincidence analysis (20)), but was so far limited to *in vitro* systems such as purified proteins (21) or DNA oligonucleotides (20) labeled with organic dyes. We performed TRICS on data obtained on cells co-expressing PC subunits PA-mEYFP, PB1-mEGFP, and PB2-mCherry2 or cells co-expressing free mEGFP, mEYFP, and mCherry, as a negative triple-correlation control. To evaluate ternary complex formation, we quantified the relative triple-correlation (rel.3C., see Materials and Methods) for both samples from the amplitudes of the ACFs and triple-correlation functions (3CFs). Fig.7A and B show representative 3CFs for the negative control and the PC sample, respectively. For the negative control, we obtained rel.3C. values fluctuating around zero (Fig.7C), rel.3C.=-0.02±0.54 (mean±SD, n=49 cells). In contrast, significantly higher, positive rel.3C. values were obtained for the polymerase samples, rel.3C.=0.43±0.38 (mean±SD, n=53 cells). The detection of ternary complexes is limited by non-fluorescent FPs, i.e. only a fraction of ternary complexes present in a sample will emit coincident signals for all three FP species. In addition, imperfect overlap of the detection volumes for each channel will further reduce the fraction of ternary complexes that can be detected by TRICS. We therefore performed an approximate calculation of the expected rel.3C. value for a sample containing 100% ternary complexes assuming a *p*_*f*_ of 0.7 for each FP species and estimating the reduction due to imperfect overlap from the pair-wise rel.cc. values detected on the positive cross-correlation control (see SI, paragraph 3 for details). For a 2:2:2 stoichiometry, we obtained an estimated rel.3C. of 0.48, i.e. only slightly higher than the average value determined experimentally for IAV PCs. Thus, we estimate that around 90% of PC subunits undergo ternary complex formation in the cell nucleus when all subunits are present.

**Figure 7.**
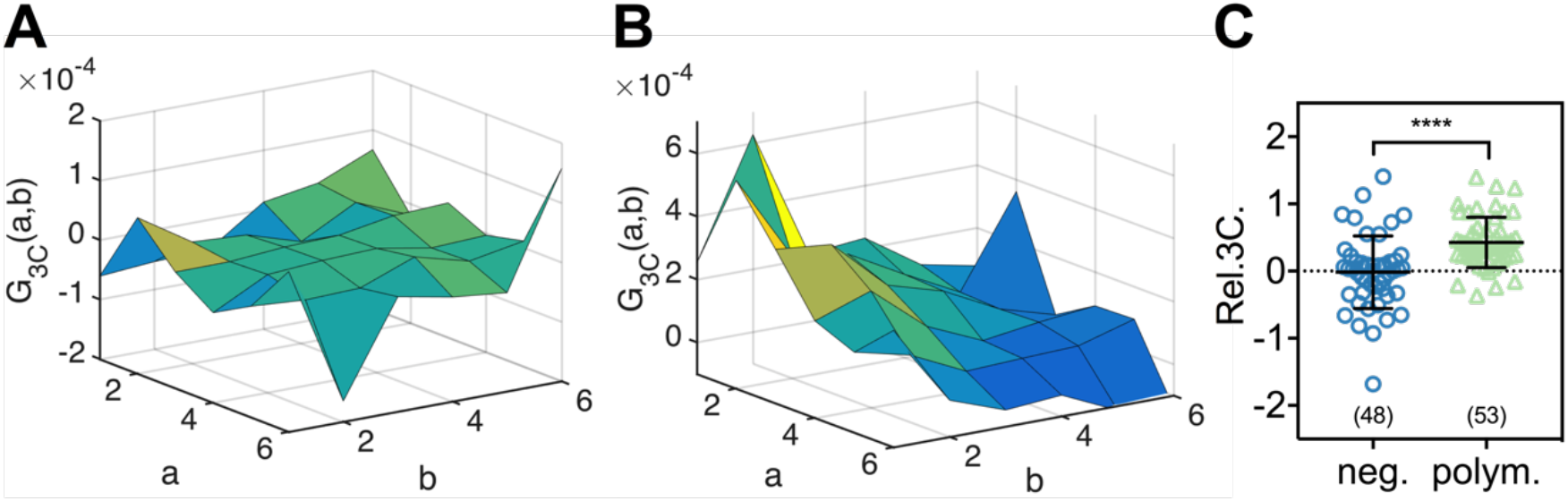
TRICS reveals the formation of ternary IAV polymerase hetero-complexes in the nucleus of A549 cells. **(A,B)** Representative 3CFs obtained from TRICS measurements on A549 cells co-expressing mEGFP, mEYFP, and mCherry2 (“neg.”) (A) or co-expressing PA-mEYFP, PB1-mEGFP, and PB2-mCherry2 (“polym.”) (B). The axes *a* and *b* indicate shifts in the x and y direction respectively, across the three detection channels, as described in the Materials and Methods. (C) Relative triple-correlation (rel.3C.) values obtained from the measurements described in (A,B). The number of cells measured is given in parentheses. Error bars represent mean±SD. Statistical significance was determined using Welch’s corrected two-tailed student’s *t*-test (\*\*\*\**P*<0.0001).

## DISCUSSION

In this work, we combine FFS techniques with spectral detection to perform multi-color studies of protein interactions and dynamics in living cells. In particular, we present SFSCS, a combination of FSCS (22) and lateral scanning FCS (11). We show that SFSCS allows cross-talk-free measurements of protein interactions and diffusion dynamics at the PM of cells and demonstrate that it is capable of detecting three or four species simultaneously. Furthermore, we extend RSICS (27) to investigate four fluorophore species and apply this approach to determine the stoichiometry of higher order protein complexes assembling in the cell nucleus. Notably, the technical approaches can be carried out on a standard confocal microscope, equipped with a spectral photon counting detector system.

In the first part, we present two-species SFSCS using a single excitation wavelength and strongly overlapping fluorophores. Compared to the conventional implementation of FCCS with two excitation lasers and two detectors, SFSCS has substantial advantages, similar to the recently presented sc-FLCCS (28). Since it requires a single excitation line and is compatible with spectrally strongly overlapping FPs, it circumvents optical limitations such as imperfect overlap of the observation volumes. This is evident from higher rel.cc. values of 70-80% measured for mEGFP and mEYFP coupled in FP hetero-oligomers compared to 45-60% observed for mEGFP and mCherry2. Rel.cc. values below 60% for green and red FPs were also observed with single-wavelength excitation (29, 38), indicating that overlap of both excitation and detection volumes (the latter requiring FPs with similar emission spectra) is required to maximize the achievable cross-correlation (29). Notably, two-species SFSCS can not only successfully discriminate between mEGFP and mEYFP, but is also applicable when using the red FPs mApple and mCherry2. These two FPs were successfully used in several FFS studies (14, 30, 39), providing the best compromise between brightness, maturation and photostability among available red FPs, which generally suffer from reduced SNR compared to FPs emitting in the green or yellow part of the optical spectrum (13, 14, 40).

In comparison to sc-FLCCS, it may be more robust to discriminate fluorophores based on spectra rather than lifetimes, which can be strongly affected by FRET (28). The emission spectra of the FPs utilized in this study did not depend on cell lines or subcellular localization (Fig.S3) and showed no (mEGFP, mEYFP) or little (mApple, mCherry2) variation with pH over a range of 5.0 to 9.2. (Fig.S10). For red FPs, specifically mApple, a red shift appeared at more acidic pH, in agreement with previous studies (41). This aspect should be considered for specific applications, e.g. RSICS in the cytoplasm containing acidic compartments such as lysosomes. Generally, spectral approaches require accurate detection of photons in each spectral bin. A previous study using the same detection system reported intrinsic cross-talk between adjacent spectral bins (30). However, since the methodology presented here is based on temporal (SFSCS) or spatial (RSICS) correlation (both excluding the correlation at zero time or spatial lag), this issue can be neglected in our analysis.

A major limitation of SFSCS is the reduced SNR of the CFs (see Fig.1) caused by the statistical filtering of the signal emitted by spectrally overlapping fluorophore species (see Fig.S4). This limitation applies to all FFS methods that discriminate different fluorophore species based on spectral (e.g. FSCS (22), RSICS (27)) or lifetime patterns (e.g. sc-FLCCS (28)). The increase in noise depends on the spectral (or lifetime) overlap of different species and is more prominent for species that completely lack “pure” channels, i.e. detection channels in which the majority of photons can be univocally assigned to a single species (27). In sc-FLCCS, this issue particularly compromises the SNR of short lifetime species (28), since photons of longer lifetime species are detected in all “short lifetime” channels at substantial relative numbers. In these conditions, sc-FLCCS could not provide reliable results with 6-fold (or higher) difference in relative protein abundance, even though the lower abundant protein was tagged with the brighter, longer lifetime FP (28). Similarly in SFSCS, CFs corresponding to mEYFP or mCherry2 were most prone to noise (Fig.1C,F), since all channels that contain, e.g., mEYFP signal also contain mEGFP signal (Fig.S3). In our experiments, cross-talk-free SFSCS analysis with two species excited with a single excitation wavelength could be performed for relative intensity levels as low as 1:10 (mEGFP/mEYFP) or 1:5 (mApple/mCherry2). In this range, SFSCS not only enabled the quantification of protein interactions via cross-correlation analysis, but also yielded correct estimates of protein diffusion dynamics and oligomerization at the PM. An improvement of the allowed relative concentration range can be achieved by using brighter or more photostable fluorophores, e.g. organic dyes, compensating for reduced SNR due to statistical filtering. Alternatively, FP tags could be selected based on proteins oligomerization state, e.g. monomeric proteins exhibiting low molecular brightness should be tagged with fluorophores that are less prone to noise. It should be noted that the limitation of reduced SNR due to excess signal from another species also applies to conventional dual-color FCCS: bleed-through from green to red channels can be corrected on average, but reduces the SNR in red channels (42), unless more sophisticated schemes such as pulsed interleaved excitation (43, 44) are applied.

Having demonstrated that two-species SFSCS is feasible with a single excitation wavelength in the green (mEGFP, mEYFP) or red (mApple, mCherry2) part of the visible spectrum, we finally implemented three- and four-species SFSCS as well as four-species RSICS. Three- and four-species SFSCS/ RSICS do not further compromise the SNR of CFs detected for mEGFP and mEYFP, but may additionally reduce the SNR of CFs corresponding to red FPs (in particular when mEGFP and/ or mEYFP concentration is much higher than that of red FPs). For this reason, three- and four-species analysis was restricted to cells with relative average intensity levels of 1:5 or less between species with adjacent emission spectra. In this range, the increase in noise due to statistical filtering was moderate and benefited from the fairly large spectral separation of green/yellow and red emission. In addition, the higher molecular brightness of mApple (compared to mCherry2) compensated for the larger overlap of this FP with the tail of mEYFP emission. The excitation power for red FPs was generally limited by the lower photostability of mApple, which could be responsible for consistently lower rel.cc. values of mEGFP or mEYFP with mApple than with mCherry2. Nevertheless, four-species SFSCS and RSICS could successfully resolve different combinations of strongly overlapping FP hetero-oligomers, e.g. a mixture of mEGFP-mCherry2 and mEYFP-mApple hetero-dimers, at the PM or in the cytoplasm of cells. To explore the interaction of four different FP-tagged proteins, four-species FFS may substantially reduce the experimental effort, because all pair-wise interactions can be quantified in a single measurement (instead of six separate conventional two-species FCCS measurements). Yet, weak interaction of proteins, i.e. a low amount of hetero-complexes compared to a high amount of unbound proteins, may not be detectable, due to the large noise of the CCF in this case. The SNR might be further compromised by slow FP maturation or dark FP states, limiting the amount of complexes that simultaneously emit fluorescence of all bound FP species (14). Ultimately, the mentioned limitations currently restrict SFSCS and RSICS to four FP species. The approaches would thus strongly benefit from a multiparametric analysis. For instance, combining spectral and lifetime detection schemes would provide additional contrast for photons detected in the same spectral bin. This improvement could expand the range of detectable relative concentrations or might allow further multiplexing of FFS.

Conventional two-color scanning FCCS has been previously applied to quantify receptor-ligand interactions in living zebrafish embryos (12) and CRISPR/Cas9 edited cell lines to study such interactions at endogenous protein level (45). SFSCS is thus directly applicable in the complex environment of living multicellular organisms. In this context, spectral information could be further exploited to separate low signal levels of endogenously expressed, fluorescently tagged proteins from autofluorescence background.

As a first biological application of SFSCS, we investigated the interaction of IAV matrix protein M2 with two cellular host factors: the tetraspanin CD9 and the autophagosome protein LC3. We observed strong association of LC3 with M2, and consequent recruitment of LC3 to the PM (Fig.S7), in agreement with previous *in vitro* and localization studies (34). Interestingly, molecular brightness analysis reported oligomerization (dimers to tetramers) of M2, but indicated a monomeric state of LC3 at the PM, i.e. binding of LC3 to M2 in an apparent stoichiometry of 1:2 to 1:4. However, each M2 monomer provides a binding site for LC3 in the cytoplasmic tail (46). A more detailed analysis of our data showed that in the analyzed cells (i.e. cells showing clear membrane recruitment of LC3, Fig.S7A,B), the PM concentration of LC3 was on average only 30% compared to that of M2 (Fig.S7C), although both proteins were expressed in comparable amounts in the sample in general. This suggests that not all potential binding sites in the cytoplasmic tail of M2 may be available to fluorescently tagged LC3, either due to binding of endogenous LC3, other cellular host factors, or steric hindrance. In contrast to the case of LC3, we did not detect significant binding of M2 with the tetraspanin CD9, a protein that was previously shown to be incorporated into IAV virions and supposedly plays a functional role during the infection process (47, 48). In future studies, the approach presented here may be used to further elucidate the complex interaction network of viral proteins, e.g. matrix protein 1 (M1) (49), M2, HA, and neuraminidase, cellular host factors, and PM lipids (50) during the assembly process of IAV at the PM of living cells (51).

Finally, we demonstrated that RSICS allows the quantification of the stoichiometry of higher order molecular complexes, based on molecular brightness analysis for each FP species. As example of an application in a biological context, we determined the stoichiometry of the IAV PC. Our data provide strong evidence for a 2:2:2 stoichiometry of the PC subunits PA, PB1 and PB2, i.e. dimerization of hetero-trimeric PCs. Such interactions were previously proposed based on experiments in solution using X-ray crystallography and cryo-electron microscopy (52), co-immunoprecipitation assays (53, 54), as well as single channel brightness analysis of FCCS data (for the PA subunit) (36). Intermolecular interactions in the PC are hypothesized to be required for the initiation of vRNA synthesis during replication of the viral genome (52, 55).

The results presented here provide the first quantification of these interactions in living cells, and a direct estimate of the stoichiometry of PCs in the cell nucleus. The formation of ternary PC complexes in these samples could be extrapolated from the observed high rel.cc. values for all three pair combinations, indicating very low amounts of unbound PA, PB1 or PB2 and higher order interactions (see SI, paragraph 1 for additional details). Furthermore, this observation could also be directly confirmed by performing, for the first time in living cells, a triple correlation analysis (TRICS), indicating the presence of a considerable amount of PA-PB1-PB2 complexes. It is worth noting though that the detection of coincident triple fluctuations is prone to considerable noise and thus still limited to molecular complexes present at low concentration and characterized by high molecular brightness for each fluorophore species (21, 56).

Of note, the RSICS approach presented here provides for the first time simultaneous information on molecular interactions, molecular brightness (and thus stoichiometry), diffusion dynamics, and concentration for all three complex subunits. This specific feature opens the possibility of a more in-depth analysis. For example, it is possible to quantify the relative cross-correlation of two subunits, e.g. PA and PB2, as a function of the relative concentration of the third subunit, e.g. PB1 (Fig.S9A). Similarly, molecular brightness and diffusion coefficients can be analyzed as a function of the abundance of each subunit (Fig.S9B,C). With this approach, it is therefore possible to distinguish specific molecular mechanism, e.g. inefficient PA-PB2 interactions in the presence of low PB1 concentration or efficient hetero-trimer dimerization when all subunits are present at similar concentrations. The employed experimental scheme offers a powerful tool for future studies, exploring, for example, interaction of the PC with cellular host factors or the development of inhibitors that could interfere with the assembly process of the complex, as a promising therapeutic target for antiviral drugs (57).

In conclusion, we present here three-species and, for the first time, four-species measurements of protein interactions and diffusion dynamics in living cells. This is achieved by combining and extending existing FFS techniques with spectrally resolved detection. The presented approaches provide a powerful toolbox to investigate complex protein interaction networks in living cells and organisms.

## MATERIALS AND METHODS

### Cell culture and sample preparation

Human embryonic kidney (HEK) cells from the 293T line (purchased from ATCC^®^, Manassas, VA, USA, CRL-3216TM) and human epithelial lung cells A549 (ATCC^®^, CCL-185TM) were cultured in Dulbecco’s modified Eagle medium (DMEM) with the addition of fetal bovine serum (10%) and L-Glutamine (2 mM). Cells were passaged every 3–5 days, no more than 15 times. All solutions, buffers and media used for cell culture were purchased from PAN-Biotech (Aidenbach, Germany).

For microscopy experiments, 3 × 10^5^ (HEK) or 4 × 10^5^ (A549) cells were seeded in 35 mm #1.5 optical glass bottom dishes (CellVis, Mountain View, CA, USA) 24 h before transfection. Cells were transfected 16–24 h prior to the experiment using between 50 ng and 150 ng plasmid per dish with Turbofect (HEK) or Lipofectamin3000 (A549) according to the manufacturer’s instructions (Thermo Fisher Scientific, Waltham, MA, USA). Briefly, plasmids were incubated for 20 min with 3 μl Turbofect diluted in 50 μl serum-free medium, or 15 min with 2 μl P3000 and 2 μl Lipofectamine3000 diluted in 100 μl serum-free medium, and then added dropwise to the cells. For spectral imaging at different pH values, culture medium was exchanged with buffer containing 140 mM NaCl, 2.5 mM KCl, 1.8 mM CaCl2, 1.0 mM MgCl2, and 20 mM HEPES with pH ranging from 5.0 to 9.2.

### Plasmids and cloning

The plasmids encoding FPs linked to a myristoylated and palmitoylated peptide (mp-mEGFP, mp-mEYFP, mp-mCherry2, mp-2x-mEGFP), the full length IAV A/chicken/FPV/Rostock/ 1934 hemagglutinin (HA) construct HA-mEGFP, and the plasmids for cytosolic expression of mEGFP, mEYFP, mCherry2, 2x-mEGFP, 2x-mEYFP, 2x-mCherry2 and mCherry2-mEGFP hetero-dimers were previously described (14) and are available on Addgene.

For the cloning of all following constructs, standard PCRs with custom-designed primers were performed, followed by digestion with fast digest restriction enzymes and ligation with T4-DNA-Ligase according to the manufacturer’s instructions. All enzymes and reagents were purchased from Thermo Fisher Scientific, Waltham, MA, USA.

To obtain mp-mEGFP-mEYFP, a mp-mEGFP_pcDNA3.1+ vector was first generated by amplifying mp-mEGFP insert from the respective plasmid, and inserting it into pcDNA3.1+ vector (obtained from Thermo Fisher Scientific) by digestion with NheI and AflII. Afterwards, mEYFP was amplified from mp-mEYFP, and inserted into mp-mEGFP_pcDNA3.1+ using digestion with AflII and KpnI. To clone mp-mEYFP-(L)-mEGFP (a plasmid encoding for mp-mEYFP-mEGFP hetero-dimers with a long rigid linker peptide (L) between FPs), a mp-mEYFP-(L)_pcDNA3.1+ construct was first generated by amplifying mp-mEYFP from the respective plasmid with primers encoding for the rigid linker (see Table S2 for linker peptide sequences) and inserting it into pcDNA3.1+ vector by digestion with NheI and AflII. Then, mEGFP was inserted from mEGFP-(L)_pcDNA3.1+ (see below) by digestion with KpnI and BamHI. To generate mp-mEYFP-(L)-mCherry2-(L)-mEGFP, a mp-mEYFP-(L)-mCherry2-(L) construct was first cloned by amplifying mCherry2 from a mCherry2-C1 vector (a gift from Michael Davidson, Addgene plasmid # 54563) and inserting it into mp-mEYFP-(L)_pcDNA3.1+ by digestion with AflII and KpnI. Subsequently, mEGFP was inserted from mEGFP-(L)_pcDNA3.1+ (see below) using KpnI and BamHI restriction. The mp-mEYFP-(L)-mCherry2-(L)-mEGFP-(L)-mApple plasmid was generated by inserting an mEGFP-(L)-mApple cassette into mp-mEYFP-(L)-mCherry2-(L) by digestion with KpnI and EcoRI. The mEGFP-(L)-mApple construct was cloned beforehand by amplifying mApple from PMT-mApple (39) (a kind gift from Thorsten Wohland) and inserting it into mEGFP-(L)_pcDNA3.1+ by digestion with BamHI and EcoRI. The mEGFP-(L)_pcDNA3.1+ plasmid was obtained by amplifying mEGFP from an mEGFP-N1 vector (a gift from Michael Davidson, Addgene plasmid # 54767) (using a primer encoding a long rigid linker sequence) and inserting it into a pcDNA3.1+ vector by KpnI and BamHI restriction. The mApple_pcDNA3.1+ plasmid was generated by amplifying mApple from PMT-mApple and inserting it into pcDNA3.1+ vector by digestion with KpnI and BamHI. The mp-mApple plasmid was generated by amplifying mApple from PMT-mApple, and inserting it into mp-mCherry2 by digestion with AgeI and BsrGI. To clone mp-mCherry2-(L)-mApple, a mp-mCherry2-(L)_pcDNA3.1+ plasmid was first generated by amplifying mp-mCherry2 (using a primer encoding a long rigid linker sequence) and inserting it into pcDNA3.1+ using NheI and KpnI restriction. Afterwards, mApple was amplified from PMT-mApple and inserted into mp-mCherry2-(L)_pcDNA3.1+ by digestion with KpnI and EcoRI. The mp-mCherry2-mEGFP plasmid was cloned by inserting mp from mp-mEGFP into mCherry2-mEGFP, using digestion with NheI und AgeI. The plasmids mEYFP-(L)-mApple, mEYFP-(L)-mCherry2-(L)-mEGFP and mEYFP-(L)-mCherry2-(L)-mEGFP-(L)-mApple were generated by amplifying the respective insert from mp-mEYFP-(L)-mApple, mp-mEYFP-(L)-mCherry2-(L)-mEGFP or mp-mEYFP-(L)-mCherry2-(L)-mEGFP-(L)-mApple and inserting it into pcDNA3.1+ vector by digestion with NheI and XbaI. The mp-mEYFP-(L)-mApple construct was cloned beforehand by inserting mApple from mEGFP-(L)-mApple into mp-mEYFP-(L)_pcDNA3.1+ using restriction by BamHI and EcoRI.

The CD9-mEGFP plasmid was cloned by amplifying CD9 from pCMV3-CD9 (obtained from SinoBiological #HG11029-UT, encoding human CD9) and inserting into an mEGFP-C1 vector using restriction by HindIII and BamHI. The LC3-mEYFP plasmid was generated by inserting mEYFP from an mEYFP-C1 vector into pmRFP-LC3 (58) (a gift from Tamotsu Yoshimori, Addgene plasmid # 21075, encoding rat LC3) using digestion with NheI and BglII. Plasmid M2-mCherry2 (mCherry2 fused to the extracellular terminus of matrix protein 2 from influenza A/chicken/FPV/Rostock/1934) was cloned by inserting mCherry2 from an mCherry2-C1 vector into mEYFP-FPV-M2 (a kind gift from Michael Veit) using restriction by AgeI and BsrGI. Plasmids encoding IAV polymerase subunits PA-mEYFP, PB1-mEGFP and PB2-mCherry2 (from influenza A/human/WSN/1933) were a kind gift from Andreas Herrmann.

The plasmids GPI-mEYFP and GPI-EGFP were a kind gift from Roland Schwarzer. GPI-mEGFP was cloned by amplifying mEGFP from an mEGFP-N1 vector and inserting it into GPI-EGFP, using digestion with AgeI and BsrGI. To generate GPI-mApple and GPI-mCherry2, mApple and mCherry2 inserts were amplified from PMT-mApple and mCherry2-C1, respectively, and inserted into GPI-mEYFP using restriction by AgeI and BsrGI.

All plasmids generated in this work will be made available on Addgene.

### Confocal microscopy system

Scanning fluorescence spectral correlation spectroscopy (SFSCS) and raster spectral image correlation spectroscopy (RSICS) were performed on a Zeiss LSM880 system (Carl Zeiss, Oberkochen, Germany) using a 40x, 1.2NA water immersion objective. For two-species measurements, samples were excited with a 488 nm Argon laser (mEGFP, mEYFP) or a 561 nm diode laser (mCherry2, mApple). For three- and four-species measurements, both laser lines were used. To split excitation and emission light, 488 nm (for two-species measurements with mEGFP and mEYFP) or 488/561 nm (for measurements including mCherry2 and mApple) dichroic mirrors were used. Fluorescence was detected in spectral channels of 8.9 nm (15 channels between 491 nm and 624 nm for two-species measurements on mEGFP, mEYFP; 14 channels between 571 nm and 695 nm for two-species measurements on mCherry2, mApple; 23 channels between 491 nm and 695 nm for three- and four-species measurements) on a 32 channel GaAsP array detector operating in photon counting mode. All measurements were performed at room temperature.

### Scanning fluorescence spectral correlation spectroscopy (SFSCS)

#### Data acquisition

For SFSCS measurements, a line scan of 256×1 pixels (pixel size 80 nm) was performed perpendicular to the PM with 403.20 μs scan time. Typically, 450,000-600,000 lines were acquired (total scan time ca. 2.5 to 4 min). Laser powers were adjusted to keep photobleaching below 50% at maximum for all species (average signal decays were ca. 10% for mEGFP, 30% for mEYFP, 40% for mApple and 20% for mCherry2). Typical excitation powers were ca. 5.6 μW (488 nm) and ca. 5.9 μW (561 nm). Spectral scanning data were exported as TIFF files (one file per three spectral channels), imported and analyzed in MATLAB (The MathWorks, Natick, MA, USA) using custom-written code.

#### Data analysis

SFSCS analysis followed the scanning FCS scheme described previously (11, 59), combined with spectral decomposition of the fluorescence signal by applying the mathematical framework of FLCS and FSCS (22, 23). Briefly, all scan lines were aligned as kymographs and divided in blocks of 1000 lines. In each block, lines were summed up column-wise and across all spectral channels, and the lateral position with maximum fluorescence was determined. This position defines the membrane position in each block and was used to align all lines to a common origin. Then, all aligned line scans were averaged over time and fitted with a Gaussian function. The pixels corresponding to the PM were defined as pixels within ± 2.5SD of the peak. In each line and spectral channel these pixels were integrated, providing membrane fluorescence time series *F*^*k*^(*t*) in each spectral channel *k*(*m* channels in total). These time series were then temporally binned with a binning factor of two and subsequently transformed into the contributions *F*_*i*_(*t*) of each fluorophore species *i* (i.e. one fluorescence time series for each species) by applying the spectral filtering algorithm presented by Benda et al. (22):

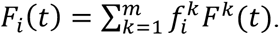

Spectral filter functions 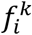 were calculated based on reference emission spectra 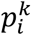 that were determined for each individual species *i* from single species measurements performed on each day, using the same acquisition settings:

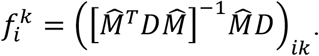

Here, 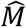 is a matrix with elements 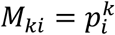 and *D* is a diagonal matrix, *D* = *diag*[1/〈*F*^*k*^(*t*)〉]. In order to correct for depletion due to photobleaching, a two-component exponential function was fitted to the fluorescence time series for each spectral species, *F*_*i*_(*t*), and a correction formula was applied (59, 60). Finally, autocorrelation functions (ACFs) and pair-wise cross-correlation functions (CCFs) of fluorescence time series of species *i* and *j* were calculated as follows, using a multiple tau algorithm:

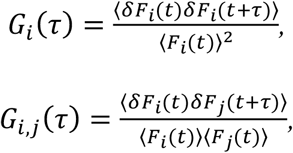

where *δF*_*i*_(*t*) = *F*_*i*_(*t*) − 〈*F*_*i*_(*t*)〉.

To avoid artefacts caused by long-term instabilities or single bright events, CFs were calculated segment-wise (10-20 segments) and then averaged. Segments showing clear distortions (typically less than 25% of all segments) were manually removed from the analysis (59). A model for two-dimensional diffusion in the membrane and Gaussian focal volume geometry (11) was fitted to all CFs:

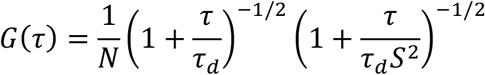

Here, the particle number *N* and thus *G*(*τ*) were restricted to positive values. In the fitting procedure, this constraint can generate low, but positive correlation amplitudes even in the absence of correlation, i.e. for CFs fluctuating around zero, due to noise. This issue causes small positive correlation amplitudes that were observed for some measurements lacking significant correlation.

To calibrate the focal volume, point FCS measurements with Alexa Fluor^®^ 488 (Thermo Fisher Scientific, Waltham, MA, USA) dissolved in water at 20 nM were performed at the same laser power. The structure parameter *S* was fixed to the average value determined in calibration measurements (typically between 4 to 8).

From the amplitudes of ACFs and CCFs, relative cross-correlation (rel.cc.) values were calculated for all cross-correlation combinations:

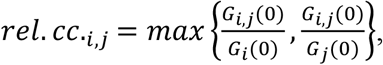

where *G*_*i,j*_(0) is the amplitude of the CCF of species *i* and *j*, and *G*_*i*_(0) the amplitude of the ACF of species *i*. The molecular brightness was calculated by dividing the mean count rate detected for each species *i* by the particle number *N*_*i*_ determined from the fit: 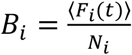. From this value, an estimate of the oligomeric state *ε*_*i*_ was determined by normalizing *B*_*i*_ by the average molecular brightness *B*_*i*,1_ of the corresponding monomeric reference, and, subsequently, by the fluorescence probability *p*_*f,i*_ for species 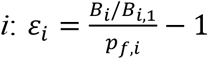, as previously derived (14). The *p*_*f*_ was previously characterized for several FPs, e.g. ca. 60% for mCherry2 (14).

The signal-to-noise ratio (SNR) of the ACFs was calculated by dividing ACF values by their variance and summing over all points of the ACF. The variance of each point of the ACF was calculated in the multiple tau algorithm (61).

To ensure statistical robustness of the SFSCS analysis and sufficient SNR, the analysis was restricted to cells expressing all fluorophore species in comparable amounts, i.e. relative average signal intensities of less than 1:10 (mEGFP/mEYFP) or 1:5 (mApple/mCherry2, three- and four-species measurements).

### Raster spectral image correlation spectroscopy (RSICS)

#### Data acquisition

RSICS measurements were performed as previously described (62). Briefly, 200-400 frames of 256×256 pixels were acquired with 50 nm pixel size (i.e. a scan area of 12.83×12.83 μm^2^ through the midplane of cells), 2.05 μs or 4.10 μs pixel dwell time, 1.23 ms or 2.46 ms line, and 314.57 ms or 629.14 ms frame time (corresponding to ca. 2 min total acquisition time per measurement). Samples were excited at ca. 5.6 μW (488 nm) and 4.6 μW (561 nm) excitation powers, respectively. Laser powers were chosen to maximize the signal emitted by each fluorophore species but keeping photobleaching below 50% at maximum for all species (average signal decays were ca. 10% for mEGFP, 15% for mEYFP, 40% for mApple and 25% for mCherry2). Typical counts per molecule were ca. 25 kHz for mEGFP (G), 15-20 kHz for mEYFP (Y), 20-30 kHz for mApple (A), and 5-10 kHz for mCherry2 (Ch2). To obtain reference emission spectra for each individual fluorophore species, four image stacks of 25 frames were acquired at the same imaging settings on single species samples on each day.

#### Data analysis

RSICS analysis followed the implementation introduced recently (27), which is based on applying the mathematical framework of FLCS and FSCS (22, 23) to RICS. Four-dimensional image stacks *I*(*x, y, t, k*) (time-lapse images acquired in *k* spectral channels) were imported in MATLAB (The MathWorks, Natick, MA, USA) from CZI image files using the Bioformats package (63) and further analyzed using custom-written code. First, average reference emission spectra were calculated for each individual fluorophore species from single-species measurements. Four-dimensional image stacks were then decomposed into three-dimensional image stacks *I*_*i*_(*x, y, t*) for each species *i* using the spectral filtering algorithm presented by Schrimpf et al. (27) (following the mathematical framework given in the SFSCS section). Cross-correlation RICS analysis was performed in the arbitrary region RICS framework (35). To this aim, a polygonal region of interest (ROI) was selected in the time- and channel-averaged image frame containing a homogeneous region in the cytoplasm (four-species measurements on FP constructs) or nucleus (three-species measurements on polymerase complex and related controls) of cells. This approach allowed excluding visible intracellular organelles or pixels in the extracellular space, but to include all pixels containing signal from the nucleus of cells. In some cells, nucleus and cytoplasm could not be clearly distinguished. In these cases, all pixels were selected and minor brightness differences between cytoplasm and nucleus, previously found to be ca. 10% (14), were neglected. Image stacks were further processed with a high-pass filter (with a moving 4-frame window) to remove slow signal variations and spatial inhomogeneities. Afterwards, RICS spatial ACFs and pair-wise CCFs were calculated for each image stack and all combinations of species *i, j* (e.g. G and Y, G and Ch2, Y and Ch2 for three species), respectively (27, 35):

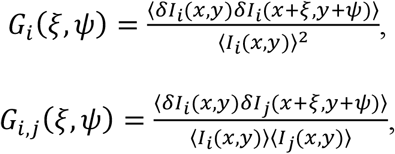

where *δI*_*i*_(*x, y*) = *I*_*i*_(*x, y*) − 〈*I*_*i*_(*x, y*)〉.

ACF amplitudes were corrected as described in (35) to account for the effect of the high-pass filter. A three-dimensional normal diffusion RICS fit model (5, 6) for Gaussian focal volume geometry (with particle number *N*, diffusion coefficient *D*, waist *ω*_n_, structure parameter *S* as free fit parameters, and fixed shape factor *γ* = 2^−3/2^ (27)) was then fitted to both, ACFs and CCFs:

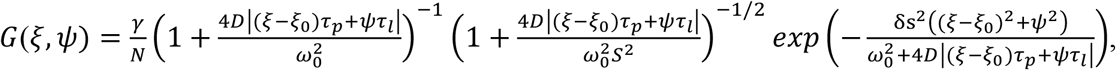

where *τ*_*p*_, *τ*_*l*_ denote the pixel dwell and line time and δs the pixel size. The free parameter *ξ*_0_ (starting value = 13 pixels) was used to determine which CCFs were too noisy (i.e. *ξ*_0_ > 4 pixels) to obtain meaningful parameters (typically in the absence of interaction). For ACF analysis, *ξ*_0_ was set to 0. To remove shot noise contributions, the correlation at zero lag time was omitted from the analysis.

From the fit amplitudes of the ACFs and CCFs, rel.cc. values were calculated:

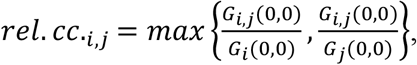

where *G*_*i,j*_(0,0) is the amplitude of the CCF of species *i* and *j*, and *G*_*i*_(0,0) the ACF amplitude of species *i*. In the case of non-meaningful convergence of the fit to the CCFs (i.e. *ξ*_0_ > 4 pixels), the rel.cc. was simply set to 0. To ensure statistical robustness of the RSICS analysis and sufficient SNR, the analysis was restricted to cells expressing all fluorophore species in comparable amounts, i.e. relative average signal intensities of less than 1:6 for all species (in all RSICS experiments). The molecular brightness of species *i* was calculated by dividing the average count rate in the ROI by the particle number determined from the fit to the ACF: 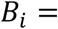 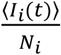. From this value, an estimate of the oligomeric state *ε*_*i*_ was determined by normalizing *B*_*i*_ by the average molecular brightness *B*_*i*,1_ of the corresponding monomeric reference, and, subsequently, by the fluorescence probability *p*_*f,i*_ for species 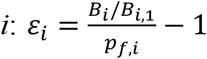, as previously derived (14). The *p*_*f*_ was calculated from the obtained molecular brightness *B*_*i,2*_ of FP homo-dimers of species 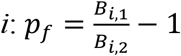 (14).

### Triple raster image correlation spectroscopy (TRICS) analysis

TRICS was performed using three-dimensional RSICS image stacks *I*_*i*_(*x, y, t*) detected for three species *i*. First, the spatial triple correlation function (3CF) was calculated:

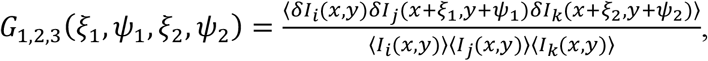

where *ξ*_1_, *ξ*_2_ denote spatial lags along lines and *ψ*_1_, *ψ*_2_ along columns of the image stacks. Contributions from *δI* triplets containing at least two intensity values from the same pixel position were not included in the calculation, in order to avoid shot-noise artefacts (since all channels are detected here by the same detector). From the resulting four-dimensional matrix, a two-dimensional representation was calculated by introducing coordinates *a, b* for the effective spatial shift between signal fluctuations evaluated for the two species combinations:

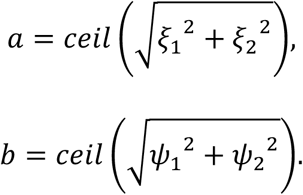

The four-dimensional triple correlation matrix was transformed into a two-dimensional representation *G*_*3C*_*(a,b)* by rounding up *a* and *b* to integer values and averaging all points with the same rounded spatial shift. For example, for a 1-pixel shift along a line in one FP channel and a 1-pixel shift along a column in the third FP channel (i.e. *ξ*_1_ = 1, *ψ*_1_ = 0, *ξ*_2_ = 0, *ψ*_2_ = 1), *a*=*b*=1. *G*_*3C*_*(1,1)* also includes in its averaged value the other seven correlation values corresponding e.g. to (*ξ*_1_ = 0, *ψ*_1_ = 1, *ξ*_2_ = 1, *ψ*_2_ = 0), (*ξ*_1_ = 1, *ψ*_1_ = 0, *ξ*_2_ = 0, *ψ*_2_ = −1) and so on. As a further example, *G*_*3C*_*(2,0)* includes and averages only the two correlation values corresponding to *ψ*_1_ = *ψ*_2_ = 0 (i.e. no shift along columns) and *ξ*_1_ = −*ξ*_2_ = ±1 (i.e. a 1-pixel shift along a line, in opposite directions for the two channels). Note that the combinations (*ψ*_1_ = *ψ*_2_ = 0, *ξ*_1_ = ±2, *ξ*_2_ = 0) and (*ψ*_1_ = *ψ*_2_ = 0, *ξ*_1_ = 0, *ξ*_2_ = ±2) would also result in *a*=2 and *b*=0, but these values were not included since they refer to a correlation between identical pixel positions (e.g. *ξ*_2_ = 0, *ψ*_2_ = 0) between two FP channels and would be influenced by shot-noise artefacts (see above).

To determine the triple correlation amplitude *G*_*3C*_*(0,0)*, the closest points (e.g. *G*_*3C*_*(1,1)*, *G*_*3C*_*(1,2)*, *G*_*3C*_*(2,1)*, *G*_*3C*_*(2,2), G*_*3C*_*(3,0)*) of the two-dimensional triple correlation were averaged, as an (slightly underestimated) approximation of the amplitude value at (0,0). Note that we chose not to include *G*_*3C*_*(2,0)* because this point is the average of only two possible spatial shift combinations, resulting in large statistical noise. Also, the point *G*_*3C*_*(0,3)* was not included since it refers to shifts along columns (i.e. the slow scanning direction) which, in turn, are characterized by a steeper decrease in amplitude. Finally, for best visualization, *G*_*3C*_ is plotted for *a* and *b* values ≥1 (see Figs. 7 and S2).

To account for reduction of the triple correlation amplitude due to the high-pass filter, an empirical correction was applied based on simulated triple correlation amplitudes with different sizes Δ*F* of the moving window (see SI, paragraph 2 and Fig.S1). Notably, applying this empirical correction to the auto- and cross-correlation amplitudes confirmed the previously introduced correction formula (see Fig.S1), 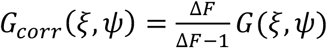 (35). The triple correlation amplitude is related to the number of triple complexes *N*_*3C*_ (20, 64):

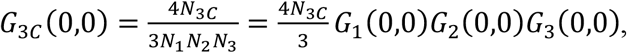

where *N*_*i*_ is the total number of proteins detected for species *i*. In analogy to the rel.cc., a relative triple correlation rel.3C. is defined, quantifying the fraction of triple complexes relative to the total number of proteins of the species that is present in the lowest concentration:

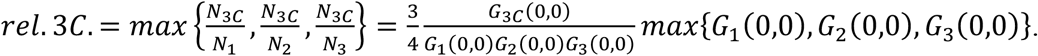

### Statistical Analyses

All data are displayed as scatter dot plots indicating mean values and SDs. Sample size is given in parentheses in each graph. Statistical significance was tested using Welch’s corrected two-tailed student’s t-test in GraphPad Prism 7.0 (GraphPad Software) and p-values are given in figure captions.

## Abbreviations

ACF: autocorrelation function
CCF: cross-correlation function
CF: correlation function
D: diffusion coefficient
F(C)CS: fluorescence (cross-) correlation spectroscopy
FFS: fluorescence fluctuation spectroscopy
(sc-)FL(C)CS: (single-color) fluorescence lifetime (cross) correlation spectroscopy
FP: fluorescent protein
FRET: fluorescence resonance energy transfer
FSCS: fluorescence spectral correlation spectroscopy
IAV: influenza A virus
M1: IAV matrix protein 1, M2, IAV matrix protein 2
mEGFP: monomeric enhanced green fluorescent protein
mEYFP: monomeric enhanced yellow fluorescent protein
mp: myristoylated and palmitoylated
HA: IAV hemagglutinin protein
PA: polymerase acidic protein
PB1/PB2: polymerase basic protein 1/2
PC: polymerase complex
pf: fluorescence probability
PM: plasma membrane
RI(C)CS: raster image (cross-) correlation spectroscopy
ROI: region of interest
RSICS: raster spectral image correlation spectroscopy
SD: standard deviation
SF(C)CS: scanning fluorescence (cross-) correlation spectroscopy
SNR: signal-to-noise ratio
TRICS: triple raster image correlation spectroscopy

## AUTHOR CONTRIBUTIONS

Research planning, V.D. and S.C; investigation, V.D. and A.P; data analysis, V.D. and S.C.; writing-original draft preparation, V.D. and S.C.; writing-review and editing, A.P.; software, V.D. and S.C.; supervision, S.C.; funding acquisition, S.C.

## ACKNOWLEDGEMENTS

This work was financed by the DFG 254850309 grant to S.C. The LSM 880 instrumentation was funded by the German Research Foundation (DFG) grant INST 336/114-1 FUGG. The authors kindly thank Madlen Luckner for providing the plasmids for PA-mEYFP, PB1-mEGFP and PB2-mCherry2 expression, Thorsten Wohland for providing the PMT-mApple plasmid and Jelle Hendrix for fruitful discussion.

