## Supplementary Info for "Multi-color fluorescence fluctuation spectroscopy in living cells via spectral detection"

### Supplementary Text

#### 1. Is pair-wise cross-correlation analysis sufficient to detect ternary interactions?

Generally, pair-wise cross-correlation analysis can only detect pair-wise interactions between fluorescently tagged protein species. To understand whether this analysis is sufficient to indicate the presence of hetero-trimeric protein complexes for the specific case reported in this work, we investigated brightness and rel.cc. data obtained by RSICS measurements of IAV PC proteins in more detail.

For all three protein species (PA-mEYFP, PB1-mEGFP, PB2-mCherry2, referred here simply as A, B and C), normalized brightness values close to the values of FP-homo-dimers were observed in this work. As a simple approximation, we assume therefore that each species, independently of its participation in hetero-complexes, is either i) exclusively dimeric or ii) present as a well-defined mixture of monomers and homo-trimers. For the latter case, the fraction of monomers ($f_{1,i}$) and trimers ($f_{3,i}$) for each species $i$ can be calculated from the average molecular brightness $\langle\varepsilon\rangle_i$:

$$f_{1,i} = \frac{1}{1+\frac{\varepsilon_{1,i}(\varepsilon_{1,i}-\langle\varepsilon\rangle_i)}{\varepsilon_{3,i}(\langle\epsilon\rangle_i-\varepsilon_{3,i})}} ,$$

$$f_{3,i} = \frac{1}{1+\frac{\varepsilon_{3,i}(\langle\varepsilon\rangle_i-\varepsilon_{3,i})}{\varepsilon_{1,i}(\varepsilon_{1,i}-\langle\varepsilon\rangle_i)}} ,$$

where $\varepsilon_{1,i}$ and $\varepsilon_{3,i}$ denote the molecular brightness of monomers and trimers, respectively.

We then calculate the maximum rel.cc. amplitudes that can be expected in the presence of optimal pair-wise interactions, while still assuming a negligible concentration of complexes containing A, B, and C.

Generally, the ACF and CCF amplitudes for multiple populations (i.e. complexes of species $i$ and $j$ with variable stoichiometry) are calculated as follows (1):

$$G_i(0,0) = \frac{\sum_k \varepsilon_{k,i}^2 c_k}{V_{\mathrm{eff}}(\sum_k \varepsilon_{k,i} c_k)^2} ,$$

$$G_{i,j}(0,0) = \frac{\sum_k \varepsilon_{k,i}\varepsilon_{k,j}c_k}{V_{\text{eff}}(\sum_k \varepsilon_{k,i}c_k)(\sum_k \varepsilon_{k,j}c_k)},$$

where $\varepsilon_{k,i}$ and $\varepsilon_{k,j}$ denote the molecular brightness of population *k* of species *i* and *j* (assumed here to be the same for all species), present at a concentration $c_k$ in the effective volume $V_{eff}$.

For the sake of simplicity, we discuss here only two simple possible scenarios for the two mixtures discussed above (i.e. each PC protein being present exclusively as homo-dimers or as a mixture of monomers and homo-trimers), in the absence of complexes containing all three PC subunits:

1) homo-dimers interacting with homo-dimers of the other species (i.e. AA-BB, AA-CC, BB-CC).

2) monomers and oligomers interacting (exclusively) with monomers or oligomers of the other species (i.e. A-B, A-C, B-C, AAA-BBB, AAA-CCC, BBB-CCC).

The two scenarios evaluated here correspond to configurations with the highest possible pair-wise correlations (in the absence of complexes containing A, B, and C), still compatible with an average oligomerization value of 2.

For the two scenarios, we calculate ACF and CCF amplitudes according to the formulas given above, assuming the same total concentration for all species and replacing the concentrations by the derived relative fractions of monomers and oligomers. For each scenario, we determine rel.cc. values from the ratio of CCF and ACF amplitudes. Finally, we extend our calculations by considering incomplete maturation of FP tags based on the fluorescence probability $p_f$. For simplicity, we assume the same $p_f$ for each FP species, in agreement with the similar $p_f$ values of ca. 60-75% observed here for mEGFP, mEYFP and mCherry2. We use a binomial model for the relative occurrence of different subpopulations in each species (2). For example, actual trimers give rise to a fraction $f_k$ of fluorescent trimers (*k=3*), dimers (*k=2*), or monomers (*k=1*) with a relative occupancy of $f_k = \binom{3}{k} p_f^k (1-p_f)^{3-k}$ and brightness $k\varepsilon_1$.

The obtained rel.cc. values for all models are given in Supplementary Table S1 for $p_f=1$ or $p_f=0.7$. For comparison, we also calculated rel.cc. values of the positive control, i.e. the maximum pair-wise rel.cc. for 1:1 stoichiometry hetero-dimers (A-B/ A-C/ B-C) or 1:1:1 stoichiometry hetero-trimers (A-B-C), resulting in values of 1 (for $p_f=1$) and 0.7 (for $p_f=0.7$). Experimentally, this control would also account for suboptimal overlap of the detection volumes for each FP combination, which we neglected here for simplicity. In the absence of ternary hetero-interactions, the determined rel.cc. values are at maximum 59% of the rel.cc. of the positive control (i.e. 0.59 for $p_f=0.7$ for scenario 1). Higher normalized values (up to 1.19, see Table S1) can be obtained only in the presence of hetero-complexes involving all three PC subunits, which we calculated for comparison for the two mixtures (i.e. AA-BB-CC, or A-B-C in mixtures with AAA-BBB-CCC) and both $p_f$ values.

Of note, in our experiments, rel.cc. values >0.7 (relative to the positive control) were observed for all pair-wise interactions between PC subunits (detected average pair-wise rel.cc. values normalized to the positive control were 0.71 for B-C, 0.97 for A-C, and 1.43 for A-B, see Fig.6D). As shown based on the different binding models, such high pair-wise rel.cc. values are only possible if ternary complexes are present. Thus, by combining molecular brightness and cross-correlation analysis, we conclude that PC proteins form a substantial amount of ternary complexes in the nucleus of cells.

**Supplementary Table S1. Relative cross-correlation values (here, same for all channel combinations) for pair-wise or ternary interactions of three-species mixtures.** Values in brackets for $p_f=0.7$ give rel.cc. values normalized to that of the positive control (i.e. the pair-wise rel.cc. for 1:1 stoichiometry).

| **Binding model** | $p_f$**=1** | $p_f$**=0.7** |
|---|---|---|
| pair-wise interactions of dimers (e.g. AA-BB, AA-CC, BB-CC) | 0.50 | 0.41 (0.59) |
| pair-wise interactions of monomers and homo-trimers (e.g. A-B, A-C, B-C, AAA-BBB, AAA-CCC, BBB-CCC) | 0.5 | 0.40 (0.57) |
| positive control (A-B/A-C/B-C or A-B-C) | 1.0 | 0.7 (1.0) |
| ternary interactions of dimers (e.g. AA-BB-CC) | 1.0 | 0.83 (1.19) |
| ternary interactions of monomers and trimers (e.g. A-B-C, AAA-BBB-CCC) | 1.0 | 0.80 (1.14) |

#### 2. TRICS analysis of simulated three-species RICS data

To evaluate the performance of TRICS, we analyzed first simulated RICS data. We ran Monte-Carlo simulations of three-species RICS for either i) three independently diffusing species A, B, C or ii) a hetero-trimeric species (e.g. A-B-C complexes). Two-dimensional diffusion and image acquisition were simulated with the following parameters: diffusion coefficient $D$=1 $\mu m^2/s$ (set to be the same for all species), $N$=1000 particles (for each species), waist $\omega_0$=0.2 μm, pixel size δs=0.05 μm, pixel dwell time $\tau_p$=2 μs, 256x256 pixels, 100 frames. RICS ACFs, CCFs and the TRICS 3CF were calculated. To correct for the reduction of the triple correlation due to the high-pass filter (with filter size of $\Delta F$ frames), an empirical correction was applied. To this aim, the variance and third central moment of a series of $10^5$ random numbers, sampled from a Poissonian distribution (with mean $f_0 = 10$), were calculated within windows with variable size $\Delta F$ (Fig.S1). The empirical function $f_i(\Delta F) = f_0 \left(\frac{\Delta F-1}{\Delta F}\right)^{b_i}$ was fitted to the variance ($i$=2) and third central moment ($i$=3). For the variance and third central moment, $b_2$=1.0 and $b_3$=3.4 were obtained, respectively. Thus, the reduction of variance and third central moment for a given value $\Delta F$ can be corrected using the factor $\left(\frac{\Delta F}{\Delta F-1}\right)^{b_i}$. For the variance, the determined value $b_2$ is in agreement with a previously discussed correction (3), which was used here to correct experimental ACFs and CCFs. To test whether 3CFs can be effectively corrected with the obtained $\left(\frac{\Delta F}{\Delta F-1}\right)^{b_3}$ factor, 3CFs were calculated with variable $\Delta F$ (in the range 2-16) and the amplitude values determined with or without the correction. In the latter case, fairly constant 3CF amplitudes were obtained, agreeing with the 3CF amplitude calculated without the high-pass filter (data not shown). Exemplary 3CFs for the two simulated scenarios are shown in Fig.S2. As expected, the rel.3C. values are close to 100% in the case of hetero-trimers and 0% in the case of independently diffusing monomers. The slight underestimation of the rel.3C. for hetero-trimers is likely due to the approximated interpolation of the amplitude value from only the first five points of the 3CF.

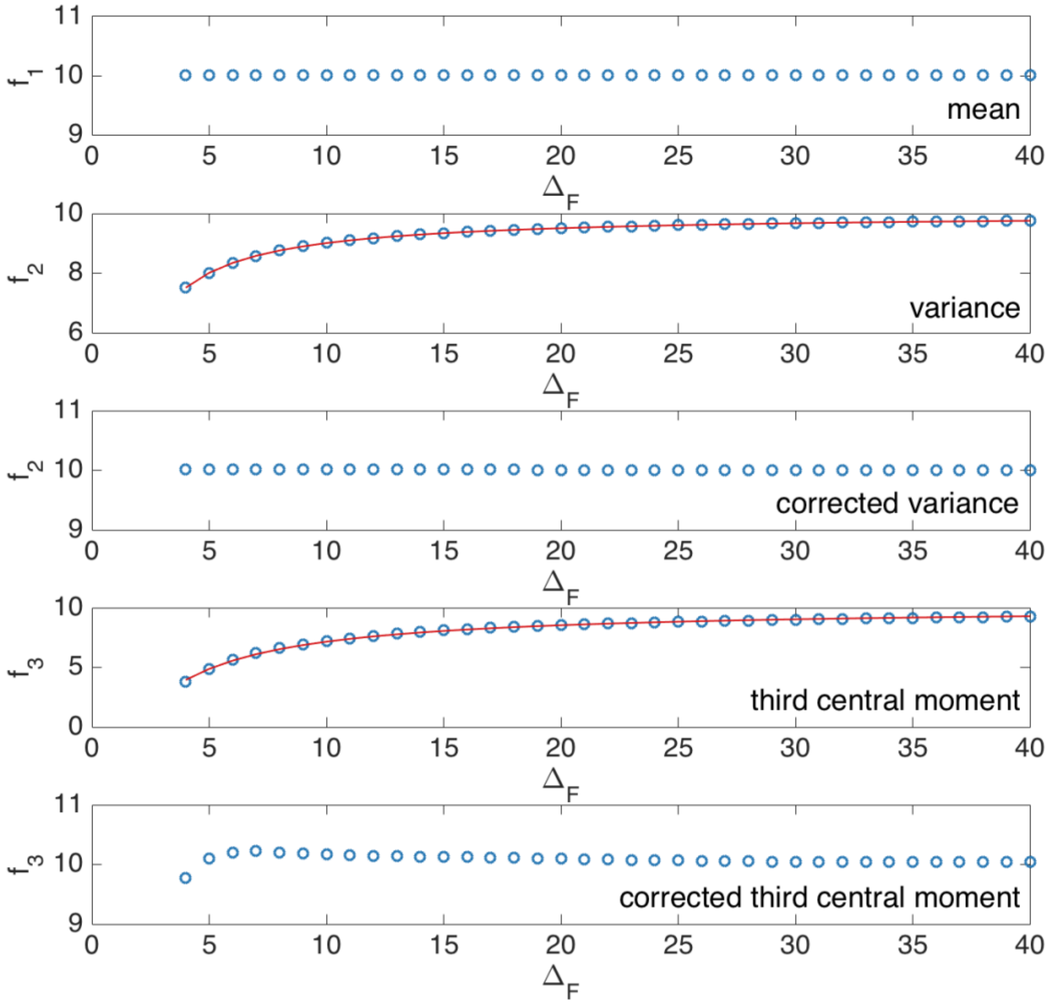


**Supplementary Figure S1. Effect of high-pass filter on calculation of variance and third central moment of random numbers sampled from a Poissonian probability distribution.** Variance ($f_2$, blue circles) and third central moment ($f_3$, blue circles) where calculated with a moving average (window size $\Delta F$) for a set of $10^5$ random numbers from a Poissonian distribution with average 10. An empirical function (red solid line) of the form $f_i(\Delta F) = f_0 \left(\frac{\Delta F - 1}{\Delta F}\right)^{b_i}$ was fitted to the variance ($f_2$) and third central moment ($f_3$), and used to correct for the undersampling effect. The corresponding values after applying the empirical correction are shown as blue circles in the panels labeled as "corrected".

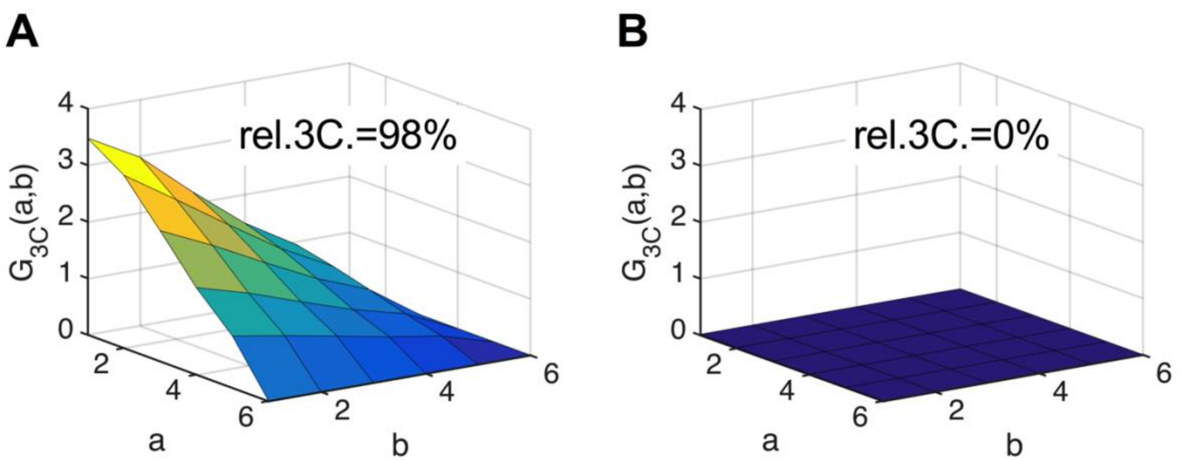


**Supplementary Figure S2. TRICS analysis of simulated three-species RICS data. (A,B)** Two-dimensional representation of the 3CF calculated for simulated TRICS data (with a 4-frame high-pass filter) for (A) ternary hetero-complexes or (B) the same number of particles per species diffusing as independent monomers. From a linear interpolation of $G_{3C}$ to *(0,0)* (using the first point $G_{3C}(1,1)$ and the average of the four points $G_{3C}(1,2)$, $G_{3C}(2,1)$, $G_{3C}(2,2)$, $G_{3C}(3,0)$) an approximate value of the 3CF amplitude was determined and corrected with the correction factor discussed in paragraph 1. The obtained value and the ACF amplitude value (also corrected for the decay induced by the high-pass filter) were used to calculate the relative triple correlation value rel.3C. (given as inset).

#### 3. Relative triple correlation for ternary complexes of fluorescently tagged proteins

The rel.3C. is a measure of the relative amount of ternary complexes in a system containing three fluorescently tagged protein species. Incomplete maturation or non-fluorescent photophysical states of FP tags will reduce the amount of detectable ternary complexes. To quantify the maximum rel.3C. that can be expected in an experiment, we calculate rel.3C. values for ternary complexes of i) *1:1:1* or ii) *2:2:2* stoichiometry, under the assumption that each fluorescent protein can be detected with a probability $p_f$. For simplicity, we assume the same $p_f$ and molecular brightness $\varepsilon$ for all three fluorophore species. Generally, the ACF and 3CF amplitudes for fully-formed ternary complexes (i.e. in absence of partially-formed complexes) of concentration *c* composed of species *1*, *2* and *3* with variable stoichiometry *l:m:n* are calculated as follows (1):

$$G_1(0,0) = \frac{c\left(\sum_{i=1}^{l}(i\varepsilon)^2\binom{l}{i}p_f^i(1-p_f)^{l-i}\right)}{V_{\mathrm{eff}}\left(c\sum_{i=1}^{l}i\varepsilon\binom{l}{i}p_f^i(1-p_f)^{l-i}\right)^2}$$ (analogously $G_2(0,0)$, $G_3(0,0)$ with upper index *m*,*n*),

$$G_{3C}(0,0) = \frac{c\left(\sum_{i=1}^{l}(i\varepsilon)^2\binom{l}{i}p_f^i(1-p_f)^{l-i}\right)\left(\sum_{j=1}^{m}(j\varepsilon)^2\binom{m}{j}p_f^j(1-p_f)^{m-j}\right)\left(\sum_{k=1}^{n}(k\varepsilon)^2\binom{n}{k}p_f^k(1-p_f)^{n-k}\right)}{V_{\mathrm{eff}}\left(c\sum_{i=1}^{l}i\varepsilon\binom{l}{i}p_f^i(1-p_f)^{l-i}\right)\left(c\sum_{j=1}^{m}j\varepsilon\binom{m}{j}p_f^j(1-p_f)^{m-j}\right)\left(c\sum_{k=1}^{n}k\varepsilon\binom{n}{k}p_f^k(1-p_f)^{n-k}\right)}.$$

From these amplitudes, the rel.3C. can be calculated (see Materials and Methods in the main manuscript). We obtain rel.3C.=$p_f^2$=0.49 (1:1:1 stoichiometry) and rel.3C.=$4p_f^2/(p_f+1)^2$≈0.68 (2:2:2 stoichiometry) for $p_f$=0.7. Due to imperfect optical overlap, experimentally detectable rel.3C. values will be lower than these values. To estimate the fraction of ternary complexes than can be detected, we compare experimental rel.cc. values obtained for all FP combinations on a positive control (FP hetero-trimers) in pair-wise cross-correlation analysis with the expected value of rel.cc.=0.7 for $p_f$=0.7 (see paragraph 1). The average rel.cc. value of 0.65 detected for mEGFP and mEYFP signal (see Fig.6D of main manuscript) was close to the expected value, hence, almost all complexes containing fluorescent mEGFP and mEYFP were detectable. On the other hand, rel.cc. values for mEGFP and mCherry2 (0.48)/ mEYFP and mCherry2 (0.53) were ca.70% of the expected value (Fig.6D). Hence, we estimate that ca. 70% of complexes carrying an mCherry2 tag and an mEGFP or mEYFP tag are detectable, due to non-optimal overlap of excitation/detection volumes. We can therefore assume that for the case of ternary complexes, ca.70% of all fully fluorescent ternary complexes that are present in the sample are optically detectable. The expected experimental rel.3C. values are thus approximately 0.34 and 0.48 for complete binding in 1:1:1 and 2:2:2 stoichiometry, respectively.

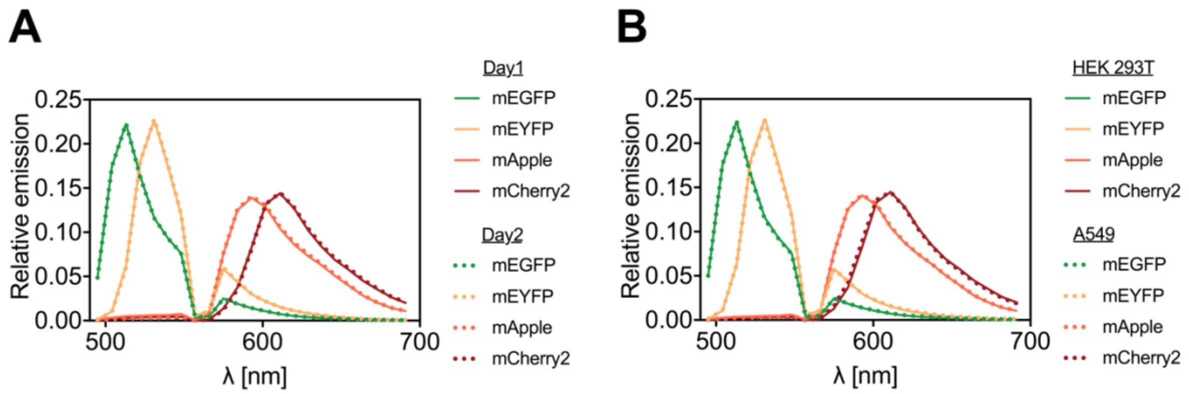


**Supplementary Figure S3. FP emission spectra. (A)** Average emission spectra of mp-mEGFP, mp-mEYFP, mp-mApple, mp-mCherry2 measured by spectral imaging (23 spectral channels from 491 nm to 695 nm) with 488 nm and 561 nm excitation on HEK 293T cells expressing each FP individually. Spectra are shown for two different days (day1: solid line, day2: dotted line) and averaged over four cells each. For each cell, 25 frames were acquired and pixels corresponding to the PM semi-manually segmented in the average image (manual selection followed by removal of pixels with intensities below 25% of the maximum pixel intensity in the selected region). **(B)** Average emission spectra measured on HEK 293T cell samples (solid line) described in (A), or on A549 cells expressing cytosolic mEGFP, mEYFP, mApple, mCherry2 (dotted line). Spectra measured on four cells each were averaged over three (HEK 293T) or two (A549) days. For A549 cells, a homogeneous ROI in the cytosol was manually selected.

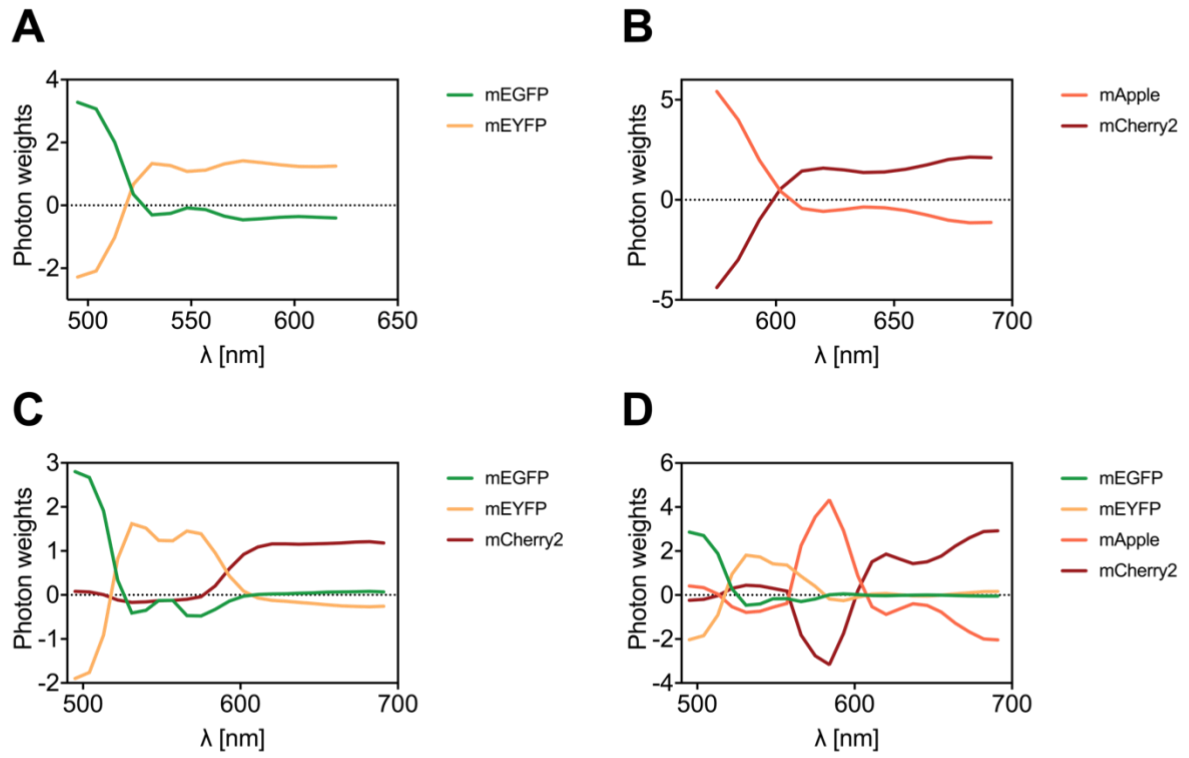


**Supplementary Figure S4. Spectral filters for two-, three-, and four-species SFSCS. (A-D)** Photon weights calculated in spectral decomposition of SFSCS data acquired on HEK 293T cells expressing mp-mEYFP-mEGFP (A), mp-mCherry2-mApple (B), mp-mEYFP-mCherry2-mEGFP (C), mp-mEYFP-mCherry2-mEGFP-mApple (D). Shown are average photon weights from five SFSCS acquisitions each.

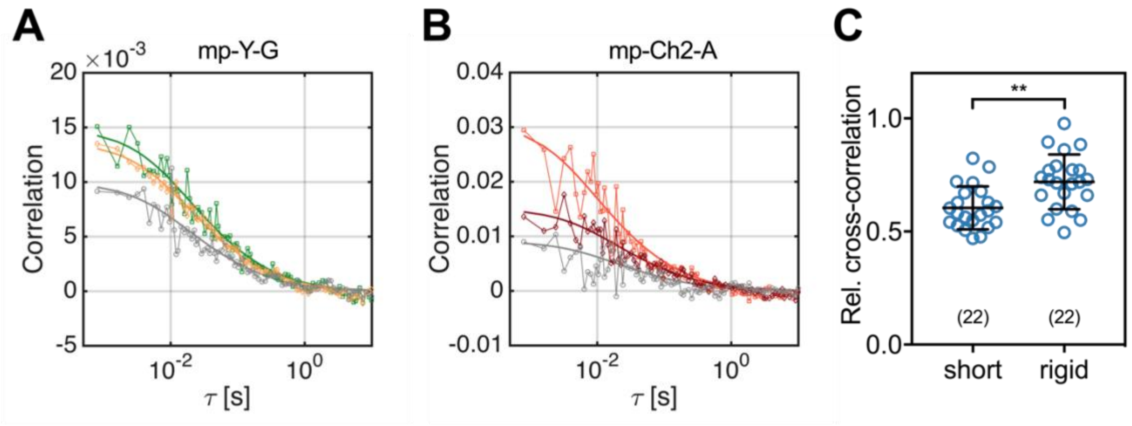


**Supplementary Figure S5. SFSCS on FP hetero-dimers. (A)** Representative CFs (green: ACF for mEGFP ("G"), yellow: ACF for mEYFP ("Y"), grey: CCF calculated between both fluorophore signals) obtained from SFSCS measurements on the PM of living HEK 293T cells expressing mp-mEYFP-mEGFP hetero-dimers. Solid thick lines show fits of a two-dimensional diffusion model to the CFs. **(B)** Representative CFs (light red: ACF for mApple ("A"), dark red: ACF for mCherry2 ("Ch2"), grey: CCF calculated between both fluorophore signals) obtained from SFSCS measurements on the PM of living HEK 293T cells expressing mp-mCherry2-mApple hetero-dimers. Solid thick lines show fits of a two-dimensional diffusion model to the CFs. **(C)** Relative cross-correlation values obtained from SFSCS measurements on HEK 293T cells expressing mp-mEYFP-mEGFP (rigid linker between the two FPs, see Table S1) or mp-mEGFP-mEYFP (short linker between the two FPs, see Table S1) hetero-dimers. Data are pooled from three independent experiments each. The number of cells measured is given in parentheses. Error bars represent mean±SD. Statistical significance was determined using Welch's corrected two-tailed student's *t*-test (***P*<0.05).

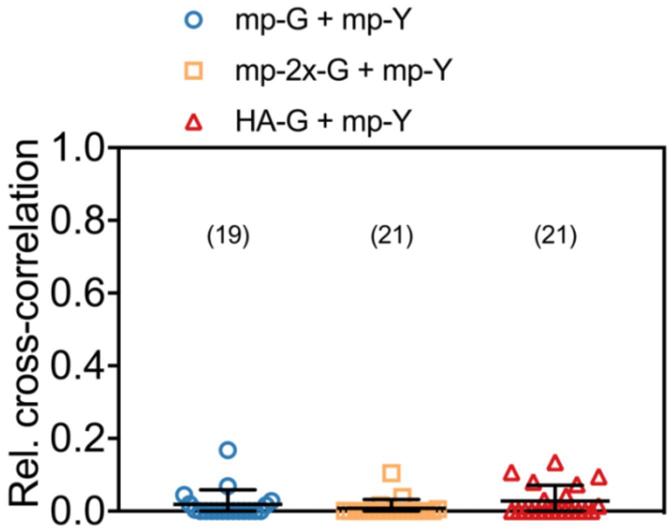


**Supplementary Figure S6. Relative cross-correlation obtained from two-species SFSCS measurements described in Fig.2 of the main text.** The number of cells measured is given in parentheses. Error bars represent mean±SD.

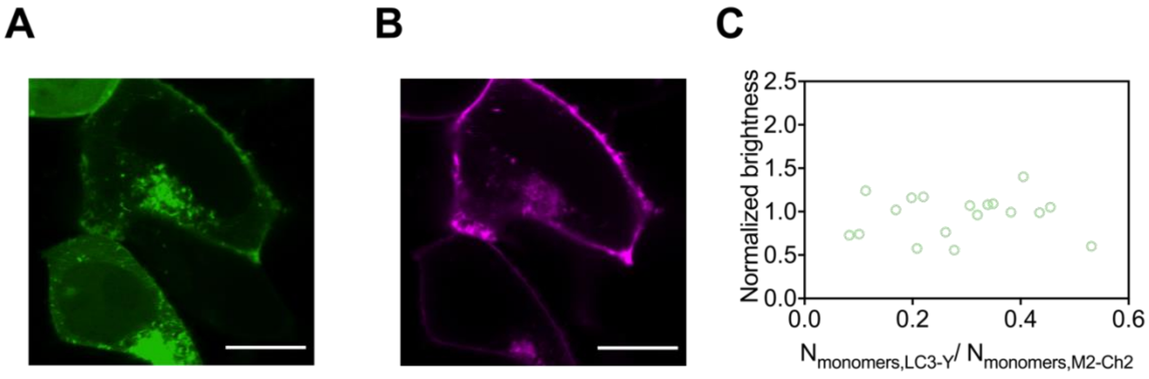


**Supplementary Figure S7. Membrane recruitment of LC3 in M2 expressing cells. (A,B)** Fluorescence images of LC3-mEYFP (A) and M2-mCherry2 (B) excited with either 488 nm (A) or 561 nm (B) excitation. LC3 is recruited to the PM in cells showing higher expression of M2 (top cell) relative to M2, but remains in the cytosol in cells expressing only low levels of M2 compared to LC3 (bottom cell). Scale bars are 10 µm. **(C)** Molecular brightness of LC3-mEYFP obtained from three-species SFSCS measurements shown in Fig.3, as a function of the ratio of LC3-mEYFP to M2-mCherry2 expression at the PM, in units of protein monomers. The number of monomers was calculated by dividing the signal detected for LC3-mEYFP/M2-mCherry2 in SFSCS measurements by the average molecular brightness detected for mEYFP and mCherry2 fluorophores in the monomeric reference sample (cells co-expressing mp-mEGFP, mp-mEYFP, and mp-mCherry2, Fig.3).

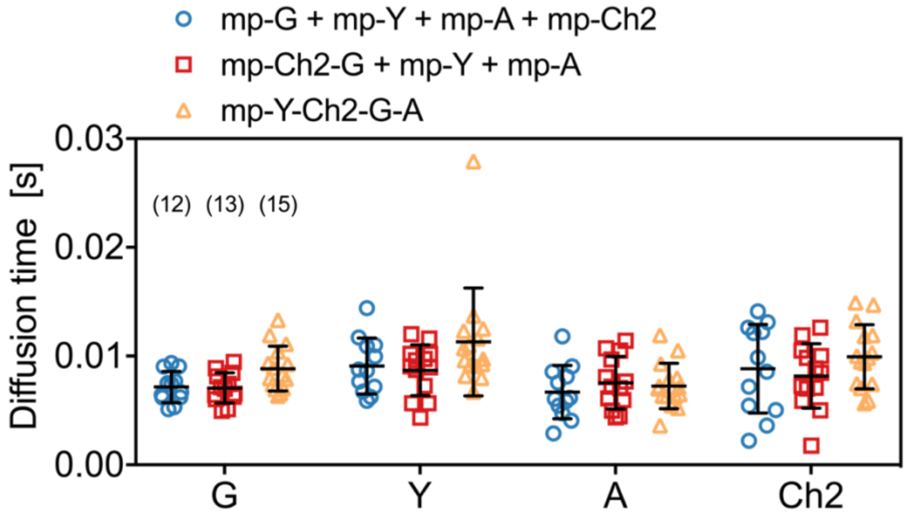


**Supplementary Figure S8. Diffusion dynamics of four-species SFSCS measurements.** Diffusion times obtained from four-species SFSCS measurements on HEK 293T cells co-expressing mp-mEGFP, mp-mEYFP, mp-mApple, and mCherry2 (blue), mp-mCherry2-mEGFP hetero-dimers, mp-mEYFP, and mp-mApple (red), or expressing mp-mEYFP-mCherry2-mEGFP-mApple hetero-tetramers (yellow). The four FP species are denoted with "G", "Y", "A", "Ch2". Data are pooled from two independent experiments. The number of cells measured is given in parentheses. Error bars represent mean±SD.

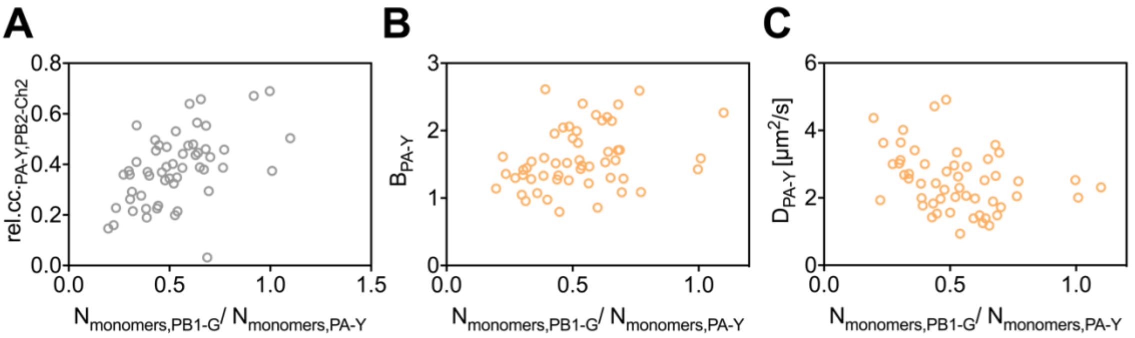


**Supplementary Figure S9. Cross-correlation and diffusion analysis for three-species RSICS measurements on IAV polymerase complex as a function of relative protein concentration. (A-C)** Relative cross-correlation for PA-mEYFP and PB2-mCherry2 (A), normalized molecular brightness (B) and diffusion coefficient (C) detected for PA-mEYFP, obtained from three-species RSICS measurements on A549 cells co-expressing PA-mEYFP, PB1-mEGFP, and PB2-mCherry2. Data are plotted as a function of the ratio of PB1-mEGFP to PA-mEYFP, in units of protein monomers, and pooled from four independent experiments (n=53 cells). The number of monomers was calculated by dividing the signal detected for PB1-mEGFP and PA-mEYFP in SFSCS measurements by the average molecular brightness detected for mEGFP and mEYFP fluorophores in the monomeric reference sample (cells co-expressing mp-mEGFP, mp-mEYFP, and mp-mCherry2, Fig.6)

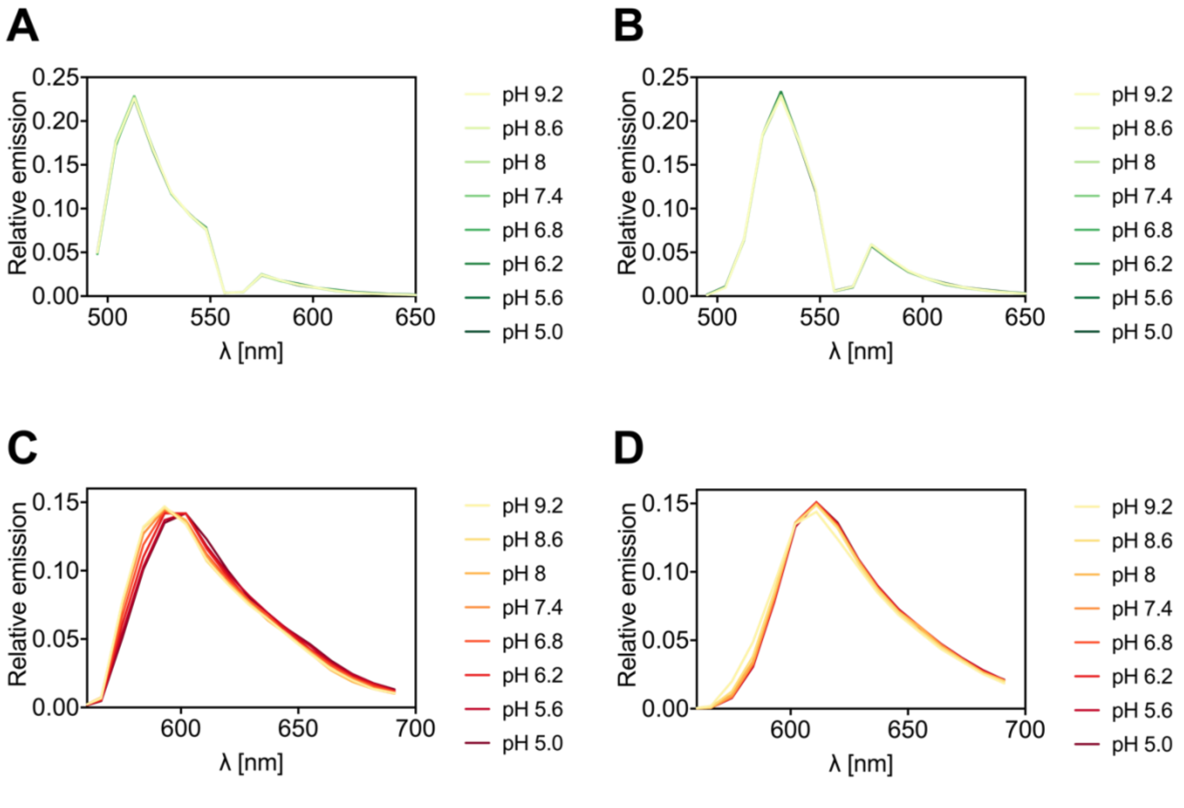


**Supplementary Figure S10. FP emission spectra at different pH values. (A-D)** Average emission spectra of GPI-mEGFP (A), GPI-mEYFP (B), GPI-mApple (C), and GPI-mCherry2 (D) measured by spectral imaging (23 spectral channels from 491 nm to 695 nm) using 488 nm and 561 nm excitation on HEK 293T cells supplemented with buffer at different pH values, ranging from pH 5.0 to pH 9.2. At each pH value, ca. 10-20 cells were imaged for five frames. To obtain average emission spectra, pixels corresponding to the PM were semi-manually segmented (manual selection followed by removal of pixels with intensities below 25% of the maximum pixel intensity in the selected region) and detected spectra averaged over all pixels and cells measured at each pH.

**Supplementary Table S2. Linker sequences of FP hetero-oligomer constructs.**

| **Plasmid** | **Linker sequence between FPs** |
|---|---|
| mp-mEGFP-mEYFP | LK |
| mp-mEYFP-mEGFP | PPAAAPPVLSLVP |
| mp-mCherry2-mEGFP | SGLRSRG |
| mp-mEYFP-mCherry2-mEGFP | 1st: PPAAAPPVLSLVP,<br>2nd: PPAAAPPVVP |
| mp-mEYFP-mCherry2-mEGFP-mApple | 1st: PPAAAPPVLSLVP,<br>2nd: PPAAAPPVVP,<br>3rd: PPAAAPPVDP |
| mp-mCherry2-mApple | PPAAAPPVVP |
| mCherry2-mEGFP | SGLRSRG |
| mEYFP-mApple | PPAAAPPVLSLVPSS |
| mEYFP-mCherry2-mEGFP | 1st: PPAAAPPVLSLVP,<br>2nd: PPAAAPPVVP |
| mEYFP-mCherry2-mEGFP-mApple | 1st: PPAAAPPVLSLVP,<br>2nd: PPAAAPPVVP,<br>3rd: PPAAAPPVDP |
